# Low-cost animal tracking using Bluetooth low energy beacons on a crowd-sourced network

**DOI:** 10.1101/2024.08.15.608178

**Authors:** DR Farine, J. Penndorf, S. Bolcato, B. Nyaguthii, LM Aplin

## Abstract

1. Animal tracking has opened the door to address many fundamental questions in ecology and conservation. While historically animals have been tracked as a means to understand their large-scale movements, such as migration, there is now a greater focus on using tracking to study movements over smaller scales, individual variation in movement, or how movements shape social network structure. With this shift in focus also comes different tracking needs, including the need to track larger numbers of individuals.
2. Tracking studies all face some technological limitations. For example, GPS and other active tracking solutions can collect fine-scale movement data, but have a high cost per tag, limiting the number of individuals that can be followed. They also have high energy costs of data acquisition and download, limiting time periods over which data can be collected. Low-energy passive (e.g. PIT) or active (e.g. reverse GPS) tags can overcome these limitations, but instead require animals to remain within a bounded study area or to come into close proximity to detectors.
3. Here we describe one solution that can overcome many current limitations by employing the massive global network of personal mobile phones as gateways for tracking animals using Bluetooth low energy (BLE) beacons. In areas with medium to high density of people, these simple-to-make beacons can provide regular updates of position over long time periods (battery life 1–3 years). We describe how to use off-the-shelf components to produce BLE beacons that weigh c. 5-6 g and cost < $7USD. Using field testing, we then show that beacons are capable of producing high-frequency tracking data that can be used to build home ranges or to detect spatio-temporal co-occurrences among individuals.
4. BLE beacons are a low cost, low energy solution for studying organisms (e.g. birds, mammals, and reptiles) living and moving in urban landscapes. Their low weight and small size makes them particularly well-suited for tracking smaller species. When combined with fixed gateways, their use can also be extended to non-urban habitats. Their high accessibility is likely to make them an attractive solution for many research projects.

## 1.0 Introduction

The ability to determine where individual animals go is central to answering many questions in behavioural ecology, conservation and management. These include identifying migration pathways (Fiedler, 2009; Alerstam & Bäckman, 2018), finding habitat refuges (Rakowski et al., 2019), identifying reproductive behaviours (Walsh et al., 2016), and understanding how animals adapt to urban landscapes (Fehlmann et al., 2024). More recently, movement patterns have also been used to study how animals move as groups (Boinski & Garber, 2000; Torney et al., 2018; Klarevas-Irby et al., 2023), make collective decisions (Strandburg-Peshkin et al., 2015; Papageorgiou et al., 2024), and form social networks (Farine et al., 2015a). Collecting individual-level movement data therefore is key to the future of research on wild animal populations (Coulson, 2020a), and as technology advances, so has the power of animal tracking methods. As a result, there is now a range of tracking tools, ranging from passive integrated transponders (PIT tags) that register in contact with specialist reader and are often deployed on hundreds of individuals (Farine et al., 2015a), to heavier and more costly GPS-based tags that can capture details of movement on a small subset of individuals (Linscott et al., 2022). Whilst each tool has its strengths, there remains a need for simple, low-cost solutions that can be deployed on to many individuals (Hebblewhite & Haydon, 2010), while also overcoming the limitations inherent with short-battery lives (e.g. having to mark many individuals simultaneously) or requiring individuals to come into/maintain close proximity to base stations or antennas.

Although there has been a plethora of tools proposed for on-board animal tracking, several main methods have dominated research to date, each with major strengths and limitations. First, radio-telemetry has involved tagging individuals with small VHF tags and following signals on foot or by drone (Rodgers, 2001; Saunders et al., 2022). This method allows real-time updates, but the number of individuals or groups that can be tracked at once is limited by researcher time. Deploying arrays of antennas to triangulate radio signals emitted from tags can automate the tracking process, making it very scalable in terms of the number of simultaneously tracked individuals (Kays et al., 2011; Fisher et al., 2020). However, these systems are relatively costly to install, complex to maintain, and are prone to shadowing by vegetation (e.g. canopy cover) or human structures (Beardsworth et al., 2022). More recently, passive integrated technology (PIT) tags have become a popular alternative, especially for smaller animals. These tags are very light-weight (<1g) and cheap to deploy per animal (<$5USD) with a small profile that allows tags to be injected or integrated into leg-rings or ear-tags (e.g. Nicolaus et al., 2008; Aplin et al., 2015b).

However, these tags are detected by changing the electromagnetic field of radio-frequency identification detection (RFID) antennas, thus requiring animals to come into very close proximity (e.g. <5cm) to the antennas. This limits the behavioural contexts for which these tags can be used, and may lead to biases in sampling, for example if individuals need to be attracted to nest-boxes or feeding stations (Kidd et al., 2015).

GPS tagging is an increasingly popular tool that can overcome many of the issues associated with radio-based tracking and PIT tags (Kays et al., 2015). Because tags use on-board capture of acquisition of the animal’s position, they can collect data from anywhere with a view of the sky. However, the costs of the individual tags means that GPS-based tracking does not easily scale up to larger populations, with studies usually limited to a small subset of individuals (Hebblewhite & Haydon, 2010a). Tracking studies that use GPS also face substantial trade-offs in terms of sampling frequency and battery life (He et al., 2022), meaning that most studies collect data at relatively low sampling frequencies (e.g. one position per hour) in order to be able to collect data over longer time frames of months to years. In some systems it is feasible to use solar panels on tags to maintain battery charge, however solar panels are easily covered up by feathers or fur, and restrict data collection to times and places with good solar recharge, introducing potential bias (Silva et al. 2017). A further challenge with GPS tagging has been the retrieval of data when the tag itself is hard to recover (Tomkiewicz et al., 2010). Common solutions have included transferring data using line of sight UHF radio, Bluetooth, mobile phone (GSM) networks, Internet of Things (e.g. LoRaWAN, or via satellite; Tomkiewicz et al., 2010; Gauld et al., 2023). Yet these all incur substantial power requirements that reduce the amount of battery available for sampling, while transmission costs can be high as more data are sampled. In some cases, there are also monetary costs involved with retrieving the data, further limiting the scale of deployment.

Here, we propose a new animal tracking solution that combines some of the advantages of passive and active tracking methods to overcome many of the limitations inherent to current technologies. Our solution builds on recently developed Bluetooth low energy (BLE) technology (Honkanen et al., 2004; Gomez et al., 2012), whereby BLE beacons emit a brief signal (called an advertisement) at set intervals that can be detected by BLE receivers (Jeon et al., 2018). In principle, this approach is similar to active RFID tags that have previously been used in a few animal studies (Ellwood et al., 2017; Chen et al., 2022), whereby tags transmit a signal that can be detected by receivers deployed throughout the environment. However, here we show how BLE beacons can additionally communicate with the most widely used BLE network—Apple’s Find My network.

The Find My network is comprised of all modern Apple devices (iPhones, watches, etc.), each of which can act as a gateway to receive signals sent by BLE beacons. When in proximity to a BLE beacon, a phone receiving the signal captures a location from its own GPS, and uploads the identity of the signalling tag and its location to Apple’s iCloud (Figure 1). Whilst initially limited to Apple Inc. devices, including Apple AirTags™, the Find My network has now been opened to third-party Bluetooth devices. As a result, it is now possible to construct custom-made beacons powered by open source software, for example by employing the recently developed *openhaystack* framework (Heinrich et al., 2021; Burg et al., 2022). Through Find My, it is now possible for animal trackers to tap into a massive global network of receivers, liberating users from the limitations that come from fixed base stations. This global network can then be supplemented, if necessary, with base stations at targeted locations (e.g. non-urban areas, near nesting sites, etc.).

**Figure 1.**
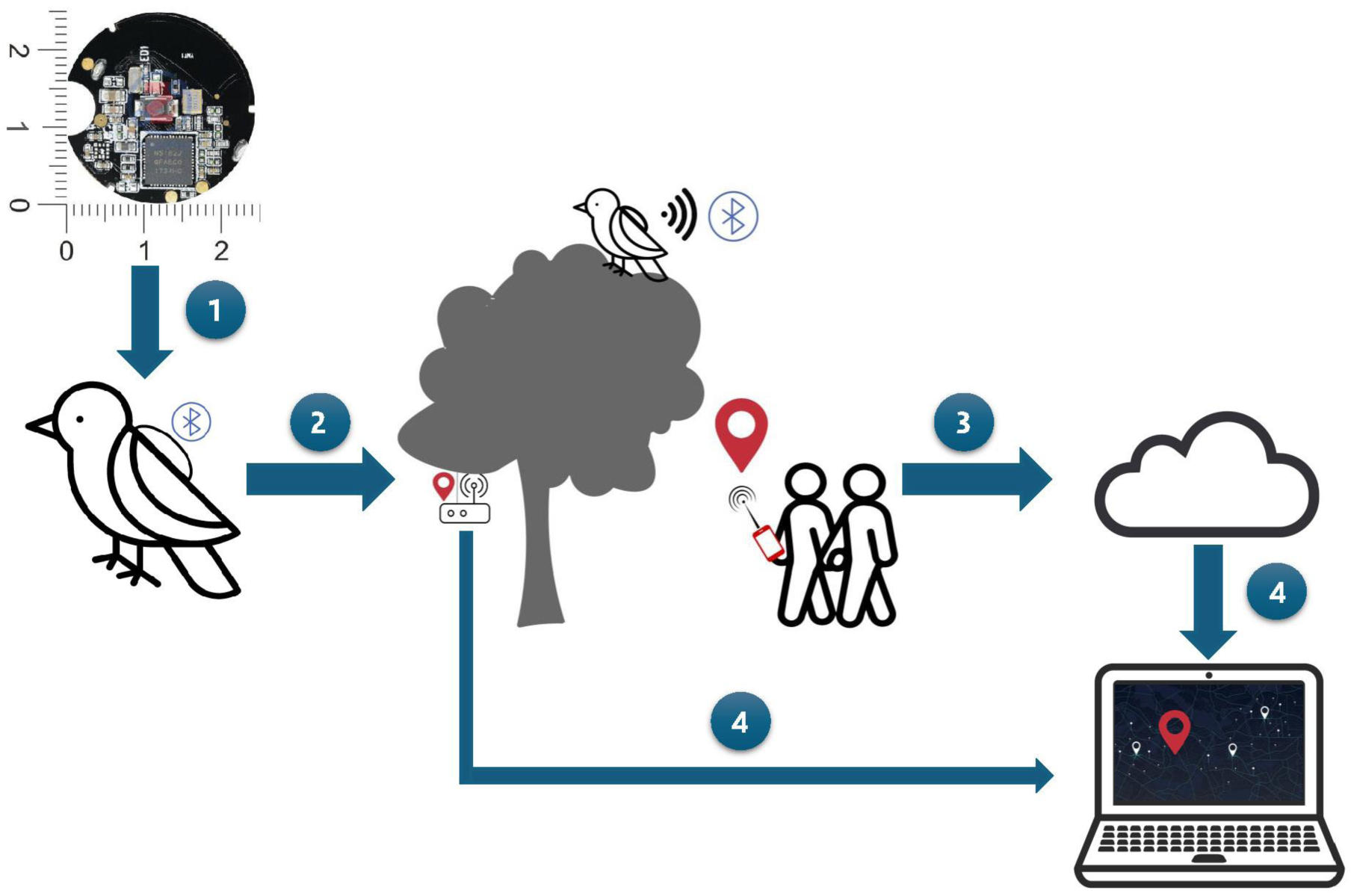
Schematic showing how data is collected from BLE beacons and how it is retrieved. (1) BLE beacons are small chips that can easily fitted to a wide range of taxa, for example by being glued onto the back of birds or fitted by use of a simple harness. (2) BLE beacons emit their unique identifier at a pre-set rate (e.g. every 2 seconds), with these beacons detectable by deployed gateways and stored onto an SD card or uploaded to a cloud, or by Apple devices. (3) When detected by an Apple device, the identity of the beacon together with a GPS point are transmitted by the device to Apple’s iCloud. (4) Data can then be downloaded from iCloud or directly from the gateways.

As we will demonstrate, in phone-rich environments such as urban landscapes, BLE beacons fitted to animals can collect data at resolutions that are comparable to GPS tags (e.g., every few minutes), while also maintaining long-term battery life (c. 3 years, depending on the advertisement interval and battery size). Finally, BLE beacons are easily built using low cost off-the-shelf products (<$10USD) with no additional installation costs, and can be built to be c. 5g. This makes BLE beacon suitable for a wide range of taxa, including birds, mammals and reptiles, and thus an exciting platform that has the potential to power many future animal-tracking studies by providing a low-cost, highly scalable option that is deployable in a range of contexts.

## 2.0 BLE beacon design

### 2.1 Bluetooth Low Energy and the Find My network

Bluetooth low energy (BLE) beacons are simple devices that emit a signal at set intervals over 2.4 GHz radio frequencies, with that signal consisting of a unique public key. BLE works in a very similar manner to classic Bluetooth, but gives a similar communication range of up to 150m with line-of-sight while operating with very low power consumption (Gomez et al., 2012b). These signals can be received by a large range of devices and BLE is now incorporated in various hardware ranging from fitness watches to security systems. Famously, and most relevant to this case, it is utilised by Apple Inc. as part of its Find My network and application. Here, when a beacon’s signal (specifically, its public key) is received by an Apple device connected to the cloud, the device fetches its current location via its location services (e.g. GPS), encrypts the location using the public key, and uploads the encrypted data to the Apple servers (Heinrich et al., 2021). This action is performed automatically by devices such as iPhones that are connected to iCloud. Apple Inc. then temporarily stores this information on its cloud (for 48 hours). During this time, the data can be queried and downloaded using the same public key, and the location information along with the estimated location error can be decrypted using a private key that is matched to the public key (i.e. known only to whoever deployed the beacon).

It is important to note that no information about the phones are recorded by the Find My network — phones simply act as data gateways without any trace of the signal having gone through the phone or of the phone’s identity in the location data that is made available. Thus, from an animal tracking perspective, there should be no privacy concerns either from the research side or for that of the general public. Security is additionally provided by requiring a private key that is matched to each public key to decrypt the location and error information (Heinrich et al., 2021).

### 2.2 Hardware

In theory, any BLE device can communicate with the Apple Find My network. The most widely used devices that are suitable for animal-borne deployments are hardware platforms based on the Nordic (www.nordicsemi.com) nRF5 chip, as these can be built into small printed circuit boards, minimising both the footprint and weight of the beacon. Conveniently, BLE beacons based on the nRF5 chip are readily available at minimal cost per unit, making them readily accessible for most research projects. The boards that we acquired for this case study weigh 1.48g without battery or casing. Boards are usually built to take a standard CR2032 battery weighing 2.9g, but can also be fitted with smaller, lighter batteries (albeit resulting in shorter battery life).

### 2.3 BLE beacon management

Because each BLE beacon requires a unique public key, and each public key requires a matched private key, these need to be generated and programmed into the BLE beacon. We provide code (adapted from https://github.com/biemster/FindMy) that can generate a required number of key pairs in the format required for programming into each BLE beacon and for downloading the data from Find My (see Appendix 1). A nice feature of this implementation is that each key has an individual .keys file that contains the private and public keys. These can be placed into a folder that can be used when querying the Find My database, and where the filename can be changed depending on the deployment. This creates a simple structure to both manage which beacons are queried at any given time (e.g. based on the deployment status) and to easily assign names to each beacon according to the identity of the individual on which it is deployed (see Appendix 2).

### 2.4 Firmware

Once keys have been generated, each BLE beacon needs to be flashed with the appropriate firmware that contains the key and provides the parameters for the operation of the beacon (specifically the advertisement interval, or how often the beacon sends a signal). Our implementation is based on the firmware initially released at https://github.com/acalatrava/openhaystack-firmware, and full instructions are provided in Appendix 3. Our instructions include how to update the firmware with the public key and set the advertisement interval, and how to upload (i.e. flash) the firmware onto the BLE beacon.

For our testing (see section 3.0 Field testing), we used an advertisement interval of two seconds, which with a CR2032 battery should correspond to a functional operation period of approximately 1-1.5 years. Long battery life is made possible because the beacon switches into deep sleep mode between each advertisement signal, during which time it uses very little power. Increasing the advertisement interval will increase battery life, whilst advertising more often will decrease battery life. Thus, extending the advertisement interval to five seconds would extend battery life to up to 3 years.

### 2.5 Casing

Once the firmware has been installed, the beacons (2.5cm in diameter, 0.4 cm in height) need to be fitted within a waterproof casing. 3D printing can provide a low-cost solution that can be adapted to the specific needs of the study species. These can prioritise total weight in smaller species (e.g. < 5 g, when combining lightweight construction with a smaller capacity battery), or robustness and avoidance of weak points in species likely to attempt to interact with the beacon (e.g. on parrots, corvids, etc.).

Post-processing techniques can further smooth the 3D-printed casing by eliminating layer lines, therefore reducing gripability of the casing once installed on the individual. Post-processing can be done manually (e.g., sanding for polyactic acid – PLA), but some materials can be chemically smoothed, allowing the processing of several casings simultaneously. For example, prints made of acrylonitrile butadiene styrene (ABS) can be smoothed using acetone vapour, and prints made of polyvinyl butyral (PVB) can be smoothed with either isopropyl alcohol or ethanol vapour.

Beacons can then be immediately attached to feathers, fur, or skin. In some taxa, such as mammals, beacons can easily be integrated into a collar. In birds, long-term deployments could utilise harnesses similar to what is used in GPS studies (Thaxter et al., 2014).

### 2.6 Data pipeline

Once a BLE beacon is fitted with a charged battery, it will continuously emit its public key at the set advertisement interval. This signal will automatically be received and uploaded by any iPhone or other apple device connected to the cloud without needing any additional steps. The data can then be downloaded by querying the Find My database. Our implementation (adapted from https://github.com/biemster/FindMy) can be run on a Mac computer with few additional requirements (see Appendix 4). We updated the original code to simplify the management of large numbers of beacons and the automation of hourly downloads of the data into daily log files, separated by projects, on a dedicated computer. Once implemented, a computer can be programmed to automatically search through the key files within a project folder (see Appendix 2), download the data, and store that data locally in daily log files for each project separately. These log files include a timestamp, the GPS location of the device that detected the beacon, and an accuracy estimate that is based on a combination of signal strength from the beacon and the phone’s own GPS accuracy. This framework allows users to easily manage beacon deployment (i.e. by adding, renaming, and removing key files), and to collate the data on a project-by-project basis.

### 2.7 Fixed BLE gateways

The strength of the BLE beacons is that they can be detected by a massive roaming network of iPhone/Apple device users. However, this network can also be supplemented by fixed receivers. Such receivers can be commercial BLE gateways, such as those often implemented in ‘smart home’ systems. However, data-logging BLE gateways can also be constructed at low cost using arduino, raspberry pi or ESP32 development boards. In Appendix 5, we provide code and instructions on how to use a ESP32 development board as a deployable gateway (based on the Freenove ESP32-S3-WROOM). This data logger can alternate between active mode and deep sleep (e.g. in periods where no data need to be collected, such as at night) and therefore is extremely energy efficient, lasting over one week depending on battery capacity. When tested in open-field conditions, it reliably detected our BLE beacons at 150m, but this range was reduced in forested conditions to approximately 75m.

### 3.0 Field Testing and Optimisation

In order to evaluate our system, we first deployed BLE beacons in fixed (known) locations in the city of Canberra, Australia. BLE beacons were programmed to advertise their public key every two seconds and enclosed in a waterproof solid plastic casing. We assessed their performance across two metrics that we considered to represent the two most critical dimensions in the use of tracking loggers—the number of detections (GPS locations) over time and the positional error. In addition, we deployed BLE beacons in pairs to investigate the relative error between beacons, which is critical when using trackers to study social behaviour (He et al., 2022). In all cases, beacons were placed under bark in trees between knee and head height. Our deployments therefore consisted of: 1) two solitary BLE beacons deployed for one week each, and 2) two pairs of beacons in <5m proximity to each other deployed for two weeks per pair. These deployments allow us to estimate detection rates, positional error, and the error in relative position of two beacons detected within a given time span.

### 3.1 Detection rates

Over the field trial, detection rates were highest during daylight hours (7am to 8pm during the trial). Detection rate ranged from 0 to 32 per hour, again depending on time of day (Figure 2). We also found that detections were affected by overall rates of traffic (e.g., proximity to arterial roads), as well as physical barriers. For example, one beacon was positioned facing a major road, and captured substantially more points than all of the other beacons combined (68% of all data, including a matched beacon on the other side of a tree trunk facing away from the road).

**Figure 2.**
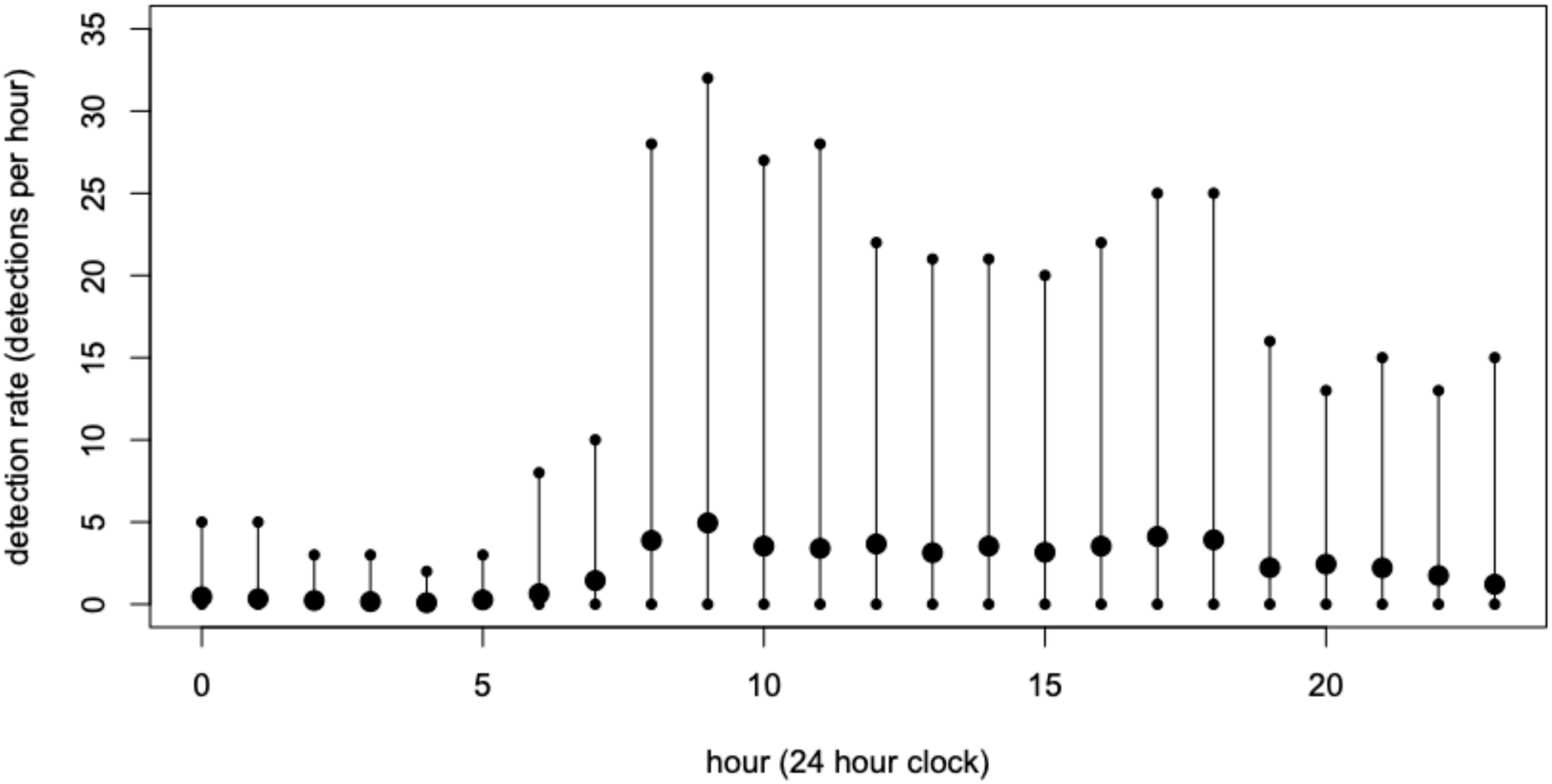
Temporal detections from BLE beacons for each hour of the day. Detection rates (number of points per hour) were variable (range of daily detection rates are shown by small points connected with lines). Deployed BLE beacons recorded between 0 points per hour to over 1 point every 2 minutes. On average (large points), beacons were rarely detected at night (typically 0 or 1 point per hour) but were detected up to 5 times per hour during the day (8am to 6pm).

### 3.2 Positional error

BLE beacons are prone to high positional error, linked to two factors: (1) GPS error on the phones on which they are detected, and (2) the distance between the phone and the beacon. This second factor means that the accuracy of the GPS on a phone does not necessarily reflect precision in the estimate of where the beacon is located. As each detection is associated with an accuracy estimate based on signal strength, it is possible to apply a threshold to detections to avoid the majority of large estimation errors. However, the relationship between the reported accuracy and the real positional error are unclear. Here, we first explored the relationship between the accuracy estimation in the data with the real positional error, representing the distance between the estimated point and the true location where the beacon was located. Positional error increased exponentially with the reported accuracy estimate (Figure 3a, note the logarithmic scale of the y axis), and this exponent increased at higher accuracy estimate (i.e. above errors of 150m or more). For a desired maximum error rate (a probability of getting an error larger than a desired error), we therefore suggest applying a threshold based on the reported accuracy estimate (Figure 3b). However, it is important to note that the data density will reduce as smaller accuracy estimates are selected.

**Figure 3.**
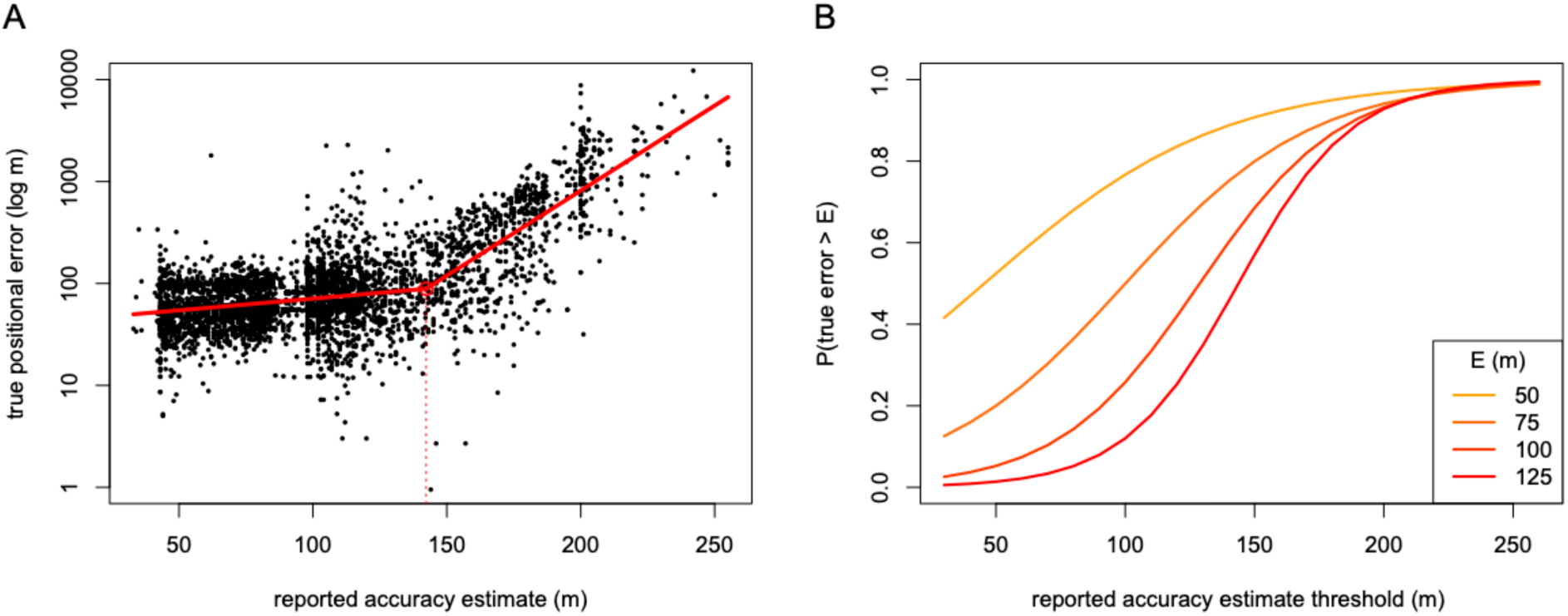
Thresholding detections based on the reported accuracy estimate can reduce the frequency of high position errors in the data. (A) The relationship between the reported accuracy estimate (error in m) reported through the Find My network and the true positional error (the distance between the reported GPS point and its real position). This relationship is exponential (note the log scale on the y-axis), with a changepoint detection algorithm suggesting that the exponent is greater for larger values of the accuracy estimates (red line). (B) The accuracy estimate allows setting a threshold to reduce the probability of getting errors that exceed a desired distance error value (E). Here we calculated, based on the real sampling data, the probability of getting an error above a desired value E when thresholding detections based on the reported accuracy estimate. For example, using a threshold of 80 m corresponds to a 15% probability of getting an error of 100 m or more.

A further finding is that positional errors were not evenly distributed in all areas (Table 1). BLE beacons that were located closest to main arterial roads had higher proportions of detections with a true positional error in excess of 150 m (21-33%). By contrast, beacons away from main roads had lower rates (5-8%). However, we note that beacons close to main arterial thoroughfares recorded substantially more data due to the higher human traffic, thus offsetting the data lost by thresholding for high error rates. In all our trials, Apple devices were rarely very close to the beacons, but rather much more likely to be on nearby paths, or in cars or houses. The distance between the beacon and the device therefore causes an intrinsic error that resulted in all beacons having a high proportion of detections with a true positional error of > 50m. Thus, we expect that our trials likely capture realistic error rates when beacons are deployed on animals.

**Table 1.** Variation in rates of true positional errors between beacons.

| Position | Road | Walkway | Road 1 | Road 2 | Residential 1 | Residential 2 |
| --- | --- | --- | --- | --- | --- | --- |
| % error > 50 m | 84 | 61 | 78 | 66 | 82 | 76 |
| % error > 100 m | 40 | 37 | 35 | 40 | 10 | 16 |
| % error > 150 m | 33 | 6 | 21 | 24 | 5 | 8 |

### 3.3 Relative position of BLE beacons detected at the same time

Simultaneous deployments of BLE beacons could be used to estimate the co-occurrence (or proximity) of animals from one-another, helping to answer a range of questions on social behaviour (Farine & Whitehead, 2015; Croft et al., 2016; Sah et al., 2018). However, the ability to generate robust estimates will rely on the relationship between the errors among beacons (He et al., 2022). Like with GPS tags, we expect that the error between beacons should be correlated, especially if they are detected by the same device. Here we explored the possibility of estimating co-occurrences of individuals within a given time window and distance by placing two pairs of BLE beacons in close proximity (< 5m). We then identified all pairs of detections that occurred within 60s of each other (note that because beacons were stationary, these results are not sensitive to the choice of window size). Finally, we quantified the proportion of positional estimates that were within a given distance threshold (Figure 4). This analysis revealed two important insights. First, the estimation of associations between individuals is not sensitive to the accuracy threshold; that is, all lines are relatively flat across the range of thresholds applied to the reported accuracy estimate. Second, we detected some differences between locations, with the estimates being substantially poorer in the pair of beacons located close to the main arterial road (Figure 4A) than in the pair placed away from roads along a suburban footpath (Figure 4B).

**Figure 4.**
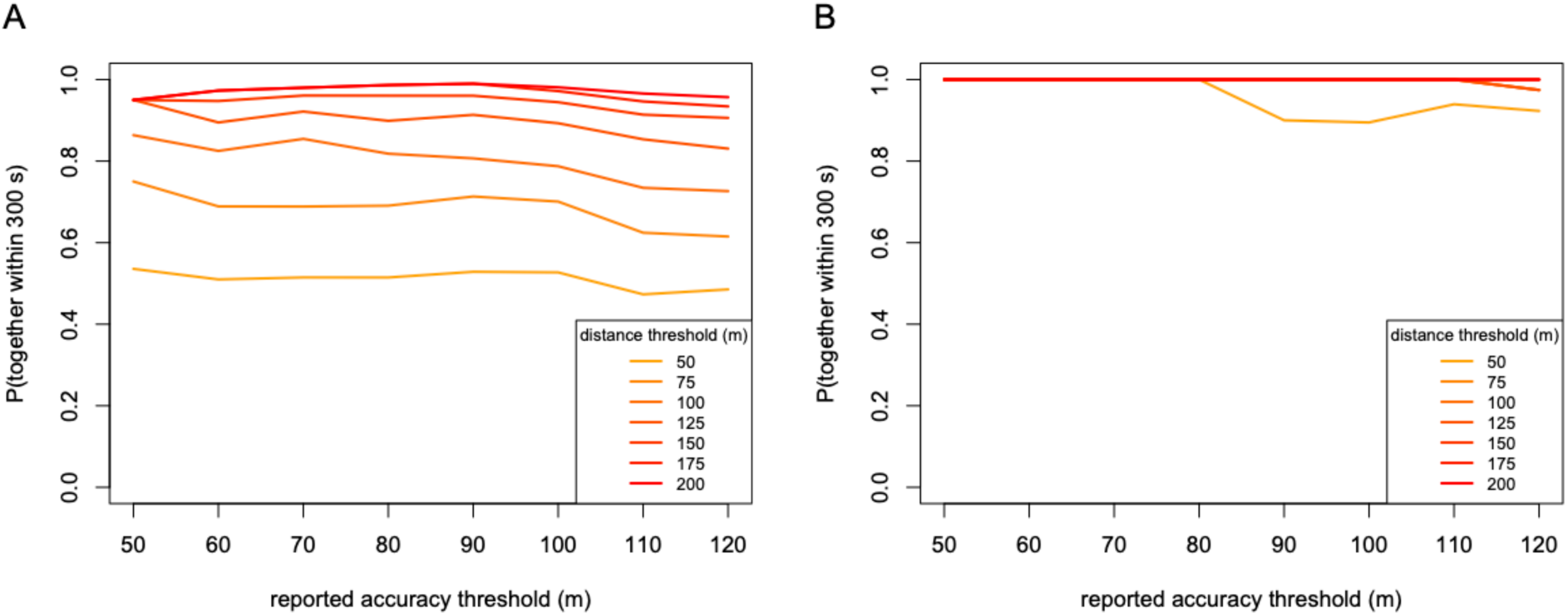
Effects of different thresholding decisions on estimates of spatiotemporal co-occurrences between pairs of BLE beacons. For each pair of beacons we quantified the proportion of detections where both beacons were detected within a 5 minute window and within a given distance threshold (line colours). (A) Proportion of detections where two beacons placed next to a road co-occurred for given accuracy and distance thresholds. (B) Proportion of detections where two beacons placed next to a footpath co-occurred for given accuracy and distance thresholds. The probability of detecting two individuals within a given distance threshold was only marginally affected by the accuracy threshold used to select which points to include in the analysis. By contrast, the rate of detections within a given distance and accuracy threshold varied between pairs. When placed next to a road (A), where there were more detections, two beacons were more often detected apart. However, overall the beacons proved to be very suitable for detecting co-occurrences of individuals.

### 4.0 Proof of Concept: detection rates in urban-living sulphur-crested cockatoos

As a proof of concept, we conducted a field test to assess detection rates from a large deployment of BLE beacons in an urban-living population of sulphur-crested cockatoos (*Cacatua galerita*). Sulphur-crested cockatoos are a large parrot (700-1200 g) that are successful urban adapters across Australia (Aplin et al. 2021, Penndorf et al. 2023), with well-known foraging adaptations that enable them to exploit human resources (Fehlmann et al. 2024, Klump et al. 2021, Klump et al. 2022). In non-breeding seasons, individuals aggregate in large communal sleeping roosts that can contain from 100 to over 1000 individual birds. Within these roosts, individuals express fission-fusion dynamics (Aplin et al. 2021), with individuals foraging up to several kilometres away before returning to the same, or a nearby, roost each evening. Here we used the large-scale deployment of BLE beacons onto sulphur-crested cockatoos to examine spatial variation in their detection rates, making use of the foraging movements of birds within and between different urban areas.

### 4.1 Deployments

We habituated sulphur-crested cockatoos across ten neighbouring roosting communities in north and south Canberra between April and June 2024, following methods from Penndorf et al. (2023). As parrots are notorious for removing attachment mechanisms commonly used in GPS-studies, such as backpack harnesses and tail-mounts (Cope et al. 2024), we glued BLE beacons on the backs of habituated individuals while the birds freely foraged within arm-reach of the experimenter. Throughout this time we deployed BLE beacons to > 200 individuals in ten distinct roosting locations, with BLE beacons lasting on the bird from 1-90 days before falling off. Each BLE beacon was fitted with a CR2016 battery and placed in a white 3D-printed casing, with the final assembled beacon weighing 6.38g, which is less than 0.91% of an individual’s body weight. All procedures were approved by the Australian National University Animal Ethics Committee (permit A2023/07).

### 4.2 Analysis

We extracted data for the time period of May 23 to June 23, obtaining detections from 178 individuals. We then partitioned detections into a 250 m by 250 m grid over the entire Canberra area, identified sequential detections for each individual that were contained within each grid, and calculated the time between detections. We restricted the daytime data 7am to 4pm to avoid over-representation of points at sleeping roosts, and only included sequential detections on the same day. We also split the data between weekdays and weekends to investigate how changes in the behaviour of people affects the detection rate over space. We then conducted our analyses at two scales. First, across all of the data, we examined the distribution of time gaps between detections (summarised for even time gaps as the BLE beacons sent an advertisement signal every two seconds). Second, we plotted the median time gap between detections over space to examine spatial variation as well as temporal changes in detection rates on weekdays and weekends. In the second analysis, we removed grid cells with fewer than 10 pairs of detections to avoid poor estimates of the median time gap between detections.

### 4.3 Results

The distribution time gap between detections fit a log-logistic distribution (Figure 5), suggesting that the time to events is generally short (the median is 110 s), but with a relatively long tail. In this case the tail is up to 9 hours, which represents birds that were detected at the roost in the morning and again at the roost in the evening, but not anywhere in between. Thus, we conclude that the extent of the tail in this distribution is over-estimated because it is caused by birds leaving a given grid cell and moving into areas where they are not detected (e.g. nature reserves). We found no different in the distribution of time gaps between detections on weekdays and weekends.

**Figure 5.**
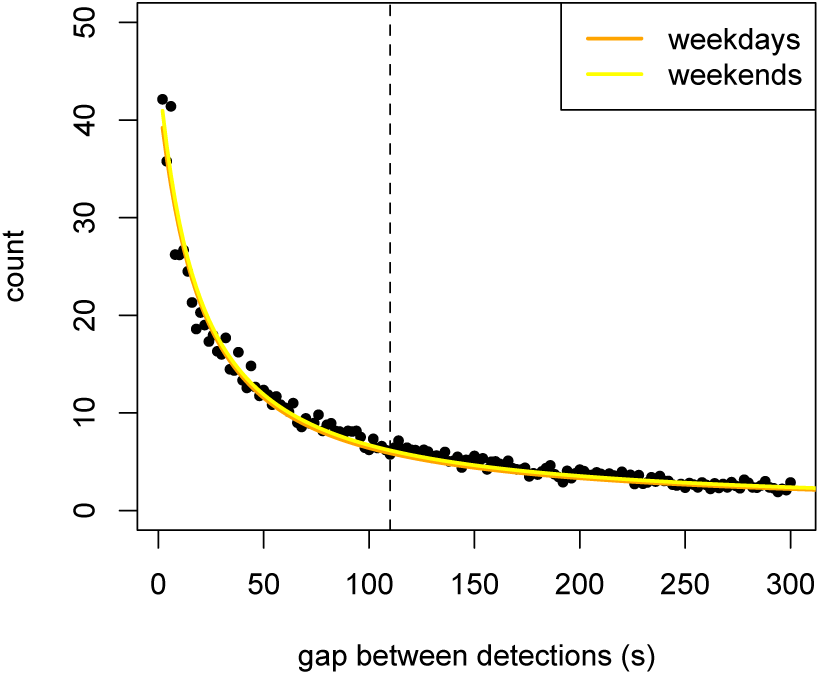
The majority of time gaps between detections are small. Points show the frequency distribution of time gaps between detections across the whole data (black points). The y-axis is the daily average count for each time gap (i.e. there were 1348 occasions in which an individual bird was detected 2 s apart over 32 days of data collection). Lines show the fit of log-logistic distributions fitted to the weekday and weekend data, showing that these are largely identical.

Within grid cells, we found that detection rates were relatively uniform (Figure 6). The range in median detection rates was from 9 s to 532 s, with a median of 132 s. This means that, on average, cockatoos were detected almost once every two minutes, with some areas having detection rates as low as once every 10 minutes. The range in detection rates did not substantially differ on weekdays and weekends (Figure 6A vs. Figure 6B). For example, the range in detection rates on weekdays was 7 s to 552 s (median 138 s) while on weekdays it was 9 s to 646 s (median 130 s). However, we did identify some areas with some predictable differences, for example a primary school and the campus of the Australian National University both had much higher detection rates during the week than during the weekend. By contrast, we did not find substantial differences in the city area.

**Figure 6.**
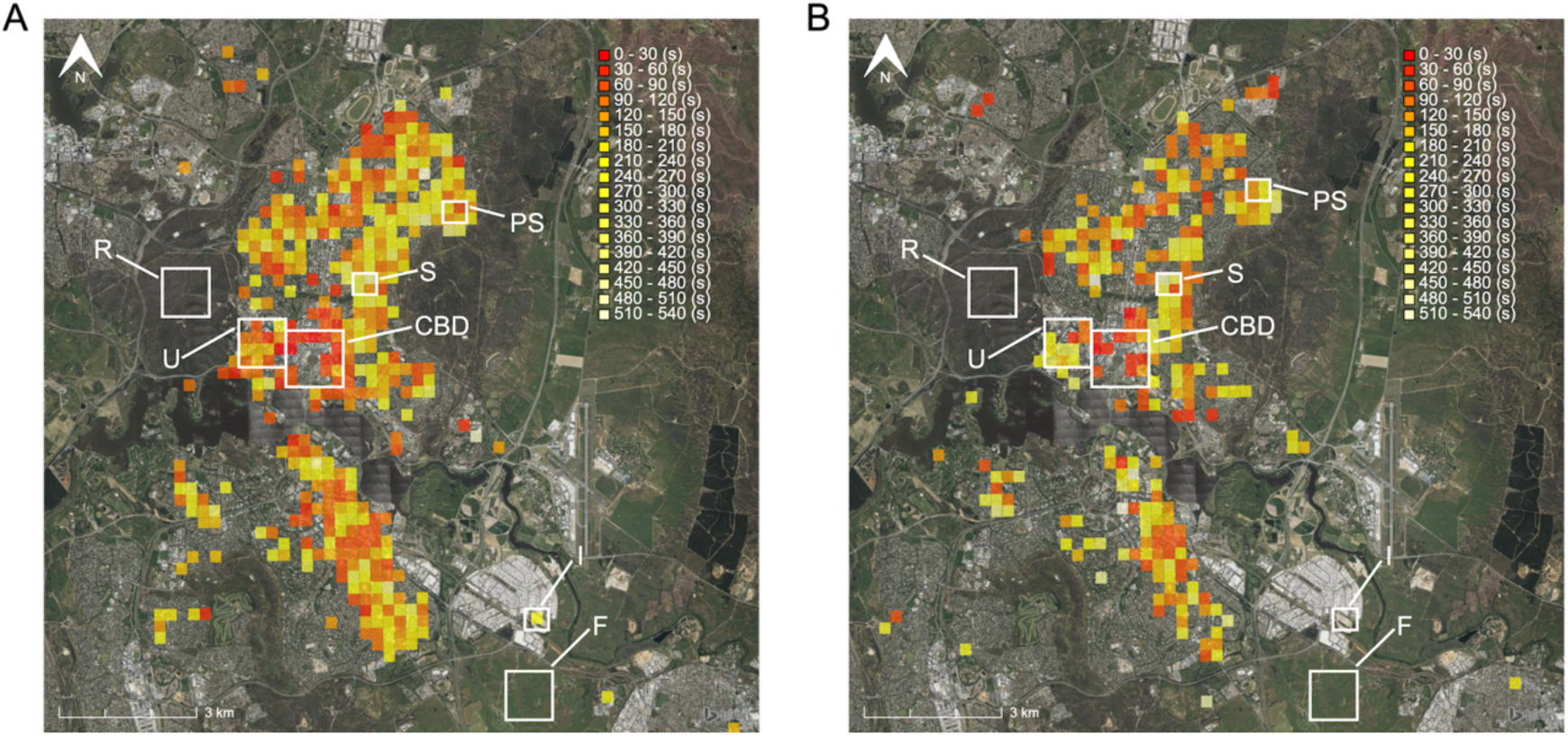
Detection rates across Canberra City for sulphur-crested cockatoos on weekdays (A) and weekends (B). Grid squares show the median time gap between detections for each grid in which cockatoos were detected at least twice consecutively. Median detection rates range from more than every 30 seconds (red) to 8-9 minutes (beige). White squares highlight different habitat types: (R) reserve, (U) university campus, (CBD) central business district, (PS) primary school, (S) typical ‘inner leafy’ suburb, (I) industrial area, (F) farmland paddocks. Variation in detection between habitat types is likely due to a combination of habitat preference by cockatoos and density of Apple devices. For example, the absence of detections in CBD is likely due to avoidance by cockatoos, while the absence of detections in reserves (R) and farmland (F) is more likely due to low density of Apple devices. Detection rates are slightly higher on weekdays on campus (U) and at school (PS), but higher on the weekend in suburbs (S). The location highlighted at (I) is a bread factory closed on weekends. In this case a stationary gateway or beacon would be required to disentangle the absence of detection on weekend versus the absence of visiting birds.

### 5.0 Proof of Concept: home range size and overlap in white-winged choughs

We conducted our second field test using white-winged choughs (*Corcorax melanorhamphos*), a group-living bird whose adult weight varies between 350–450 g (Heinsohn et al., 2000). White-winged choughs exhibit a non-territorial behaviour and form social groups of up to 20 individuals (Heinsohn et al., 2000) that remain in stable groups all year-round (Heinsohn et al., 2000). Previous studies of white-winged choughs have primarily focused on the breeding ecology, during which time groups tend to have small home ranges of up to 0.2 km2—the area immediately surrounding their nest location (Heinsohn, 1988). However, early reports (e.g. Rowley, 1978) suggest that white-winged choughs substantially increase their home range during the wintering period (up to 10km2). Here we examine the performance of Bluetooth beacons by quantifying the home range of white-winged choughs and investigate the impact of varying accuracy thresholds on the estimation of these home ranges.

### 5.1 Deployments

Between March 2024 and April 2024, we trapped, banded, and tagged eight adult choughs across four different groups with BLE beacons in the inner north region of Canberra, where the population has been monitored since 1985 (Heinsohn, 1987). The choughs were then weighed and fitted with individually numbered stainless steel metal rings from the Australian Bird and Bat Banding scheme. After processing each chough, we assembled a beacon by fitting a battery and placing the beacon in black 3D-printed casings. The BLE beacons weighed 5.7 g, representing 1.6% of an individual’s body weight. BLE beacons were attached to the birds by glueing them on the back, hence reducing the deployment time spent in comparison to attaching harnesses (which are commonly used when fitting GPS tags). We obtained the licence to trap the white-winged choughs from the Australian Capital Territory Government and the procedures were reviewed and approved by the Australian National University Animal Ethics Committee (permits A2022/35).

Our deployment focused on two birds per group from each of four adjacent social groups. This allowed us to evaluate the consistency in the estimated home range sizes across groups within the same environment, and across individuals within the same group. We expected individuals from the same group to have highly overlapping home ranges, and for individuals across groups to have adjacent and similarly sized home ranges.

### 5.2 Analysis

We conducted an estimation of the home range through the application of a weighted Automated Kernel Density Estimation method (95% wAKDE, using the R-package *ctmm*, Calabrese et al., 2016) for each individually tagged chough within the four groups (Fleming et al., 2022). The 95% home range represents the area in which the individual is estimated to occur 95% of the time. Our analysis specifically focused on fixes occurring between sunrise and sunset. Home range estimates were generated at varying distance thresholds, including the entire dataset, < 120 m, and < 80 m. The dataset was processed in R (version 4.3.3), with the subsequent plotting of maps facilitated by QGis® (version 3.30.2). Here, we report initial findings extracted from the tracking data acquired within the initial 24-day period post-deployment, enabling an evaluation of the efficacy of the Bluetooth beacons.

### 5.3 Results

The presence of clustered points in Figure 7 demonstrates high detection rates within urban regions, all of which were collected through the Find My network, thereby confirming the effectiveness of the BLE beacons in this environment. Irrespective of the accuracy threshold used, we detected clear differentiation in the space used by different groups of choughs, suggesting that the BLE beacons are capable of differentiating space use even at relatively small scales. However, the extent of the home ranges differed depending on the dataset we utilised. Initially, employing the entire dataset yielded unrealistically large and diffuse home ranges. However, when implementing a threshold based on the estimated error (accuracy), we generated home ranges that match with our expectations.

**Figure 7.**
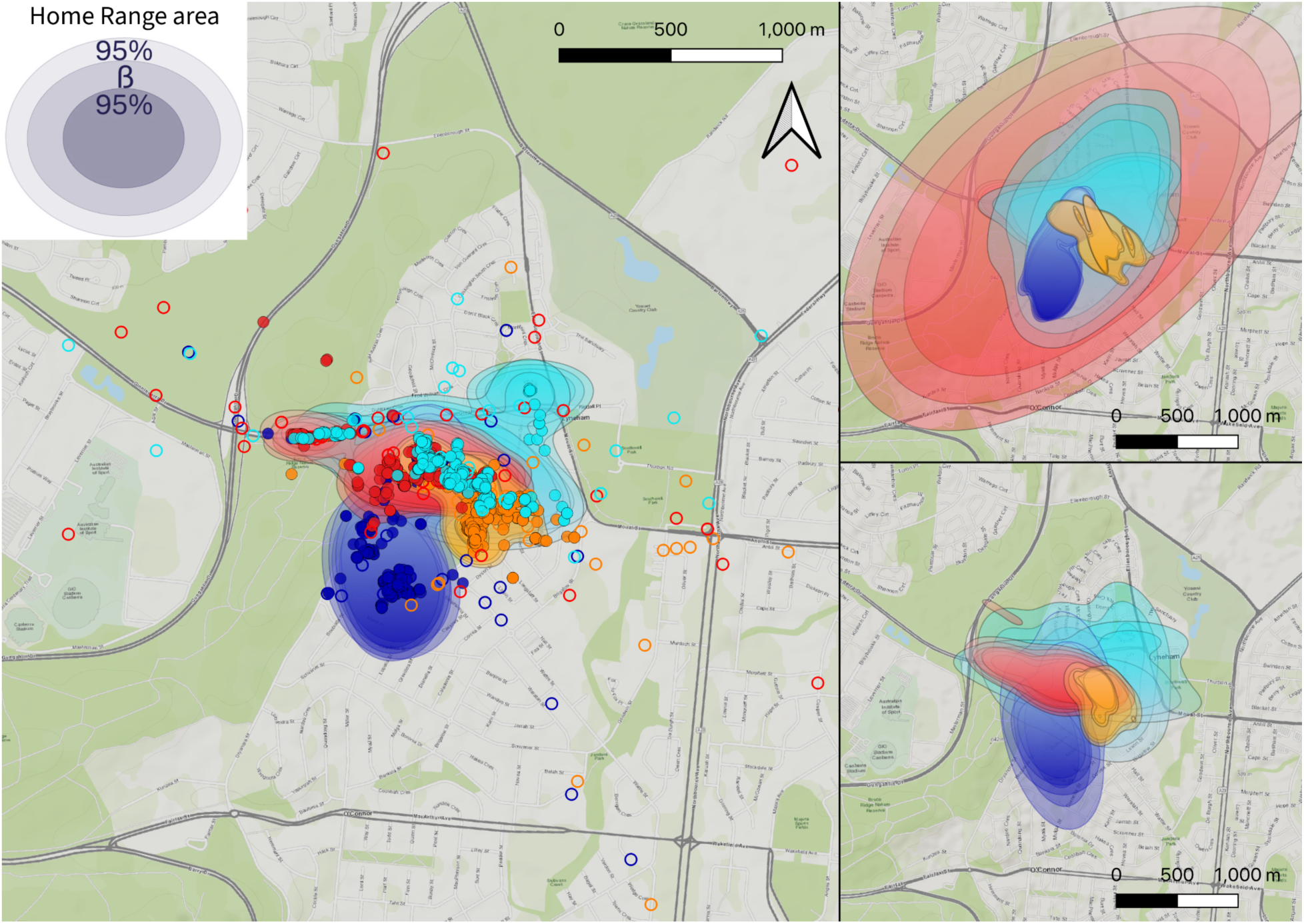
Data from beacons deployed in four groups of white-winged choughs over two months. Each colour represents the data collected from beacons fitted to two birds in the same social group (N=4 groups in total). Groups are from a study population in the inner north of Canberra, Australia. Filled points represent detections with an accuracy threshold of < 120 m (see Section 3.1), and open circles represent detections with an accuracy estimate beyond this threshold. Panels show home ranges estimated using data with a reported accuracy of less than 120 m (main panel), all of the data (top-right), and less than 80 m (bottom-right). Each home range is depicted by three lines, with the central line representing the estimate β (the area within which the individual is estimated to occur 95% of the time), and the inner and outer lines representing the 95% confidence intervals (upper and lower) of the home range estimate. Note the gaps in coverage in nature reserves (green areas), which can be addressed using gateways (see Discussion), and the effects of arterial roads in capturing a disproportionate number of points (see Section 3.1).

Interestingly, we found that a less restrictive threshold yielded clearer home ranges. More specifically, thresholding the data to < 120 m resulted in a substantially higher number of fixes, relative to a threshold of 80 m. In turn, this produced a high density of points in what is likely to be the true home range. The effect of the more extensive sample size when using the 120 m threshold is demonstrated by the more precise estimation of the home range (i.e. the 95% confidence intervals in Figure 7 are tighter) and a smaller home range size relative to the < 80 m threshold. Thus, although a 120 m threshold can include more erroneous points, the increased density of points can help ensure a greater level of resolution when estimating home ranges (as evidenced by the tighter 95% confidence intervals in Figure 7). The enhanced estimation of home ranges when using a 120 m threshold was also evident by the consistency in the estimated home range across the two members of each group. When using the < 120 m threshold, the home ranges across the two group members became almost indistinguishable. By contrast, when using < 80 m (or without using any threshold), the estimates varied substantially between individuals (e.g. the two individuals depicted in light blue).

Our observations revealed that the four white-winged chough groups maintained consistent home ranges throughout the tracking period. The mean estimated sizes of the home ranges for the four groups were (0.2 km2, 0.3 km2, 0.4 km2, 0.5 km2) across the 24-day period (when using a < 120 m threshold), aligning with the estimations in the literature for birds during the breeding period (our data overlapped with the transition from breeding to non-breeding as groups had young chicks). Therefore, our field test demonstrates that the BLE beacons can provide reliable estimates of home range size for animals living in urban environments.

### 6.0 Discussion

We provide a novel solution for tracking animals that requires minimal infrastructure or initial investment, with cheap unit costs and no ongoing data costs. Importantly, because of this low price per beacon and long battery life, Bluetooth low-energy beacons provide a potential means for projects to track larger populations than most current solutions allow. In this paper, we provide a full methodological pipeline to implement BLE beacons for animal tracking, from beacon construction to data management and download. In addition, we give guidance on how to make informed decisions on handling the data, such as how to choose a threshold based on the accuracy estimates provided with data download. We show that BLE beacons allow high rates of data collection (every few minutes, on par with state-of-the-art GPS tags) across urban landscapes. From these data, we show that BLE beacons can be used to generate accurate estimations of home ranges, and that they are suitable for studying co-occurrences among individuals. Their low price, small weight, and ease of data collection makes BLE beacons very scalable; for example, here we show data from a simultaneous deployment on > 150 individual cockatoos. This makes them particularly applicable for studies of social behaviour and extremely useful for studies spanning a wide range of organisms.

Like with all technologies, BLE beacons have trade-offs. The first major issue we have identified is error in the positional estimation. Even when thresholding based on the reported accuracy estimates, we still expect some points to have large errors. This primarily arises due to GPS error on phones combined with the distance between the phone (where the GPS point is recorded) and the beacon. While this may seem problematic, it can largely be dealt with through careful selection of research questions, study systems, and analytical tools. For example, BLE beacons may not be very suitable for studying fine-scale habitat selection or movement decisions (e.g. relative to error for GPS tags which is typically less than 10 m; He et al. 2022), but should be suitable for studying aspects like home ranges or large-scale displacements (Frair et al., 2010). This will especially be true for larger species, whose home ranges dwarf the 100 m error in positional estimates. We also note that the size of this error is comparable to reported errors from, for example, low-intensity GPS sampling and satellite tracking. Indeed, error from GPS trackers is also often substantially higher (reaching tens of meters) when the sky is obscured under canopy or in urban areas (He et al., 2022). Tracking with Argos satellite tracking tags also generally has much larger errors (in excess of 500 m; Boyd & Brightsmith, 2013). However, future work could develop solutions to reduce positional errors, which may be possible with the inclusion of additional devices on beacons, for example similar to those currently used by Apple’s AirTags (Roth et al., 2022).

A second major issue we have identified is the variation in sampling rates. We have demonstrated that within urban areas the sampling rate is likely to be adequate for almost all study aims. However, there are also large gaps in non-urban areas (e.g. reserves, farmland, or non-urban areas on weekends). This issue could be removed entirely by deploying an array of gateways (see below). However, to some extent, variation in sampling rates is a feature of most systems. For example, GPS tags can have variable sampling rates depending on the available solar or battery power. In general, estimating parameters for animal movements, such as home ranges, requires careful consideration of both error and sampling rates (Thomson et al., 2017), and sampling more individuals for longer, while using lower usage density estimates, is considered to be the best way of avoiding biased home range estimates (Börger et al., 2006). For this reason, many statistical tools that have been developed for studying, for example, home ranges can now accurately account for errors (e.g. Fleming et al., 2015). Exploration of the data for each study is likely to be quite important, for example here we identified that a threshold of < 120 m is likely to provide better estimates than using a smaller threshold, due to a greater data loss for the latter.

Thus, like with all tools, positional error and variation in detection should not limit the power of studies so long as it is accounted for in the study design (including the question and analyses).

A third, related limitation is the reliance on higher densities of mobile phones. Because of this, BLE beacons are most well-suited to studies in urban areas with a high density of Apple devices, and may struggle in more natural areas with much lower human (and thus phone) presence. Our tests were conducted in suburban areas of Canberra, a relatively low-density city in Australia of approximately 478,000 people. Detection rates in this city were good, but may be much higher in high-density areas of large cities. One solution to low detection rates is to supplement or even replace mobile phone detection with BLE gateways. Similar to arrays of RFID loggers deployed at feeders to study wild birds fitted with PIT tags (Adelman et al., 2015; Crates et al., 2016; Brandl et al., 2019; Heinen et al., 2022), BLE gateways could be used to detect tagged animals over much larger distances (approximately 100 meters for beacons versus less than 5 cms for RFID loggers).

BLE gateways are readily available to be purchased as commercial products or constructed using off-the-shelf microcontrollers; we provide instructions for programming one example in Appendix 5. Gateways can be constructed for minimal cost (e.g. < $20), and when using modern microcontrollers, they can be implemented with low power requirements, meaning that arrays could be installed over larger areas (especially when combined with solar recharge). A further advantage of this approach is that the propensity for large positional errors would be substantially reduced because gateways are stationary and at known positions. If gateways are placed at sufficiently high density, then they could even potentially be used for triangulation for finer estimates of position (Ke et al., 2018), similar to other gateway-based tracking technologies (Ripperger et al., 2020). This could be useful for estimating the relative position of individuals within a sleeping site, or their movements to or away from nesting sites. For instance, in Chen et al. (2022), gateways detecting high energy PIT tags were placed at focal roosts, nests and foraging sites of wild Eurasian jackdaws (*Coloeus monedula*), allowing tracking of movement decisions between these resources. Finally, microcontrollers could easily be interfaced with other hardware devices. For example, individuals could be detected at puzzle boxes (Aplin et al. 2015a) or other devices to facilitate identification in field experiments.

A potentially powerful approach could be to combine BLE beacons with other sensors, such as accelerometers. Accelerometers capture acceleration in three different axes, and in doing so can provide a wealth of information (we note that any such data would require an independent method of downloading the data). For example, machine learning approaches can be used to infer the behaviour of individuals at each point in time from accelerometer data (Chakravarty et al., 2019; Yu et al., 2023; Otsuka et al., 2024), which can provide critical information including how much time the individual is spending in different states (e.g. moving vs. foraging). Further, by modelling movements within high detection areas, it may be possible to extrapolate data from the accelerometer in areas without high phone coverage. Previous work has been quite successful at using dead-reckoning techniques from accelerometers to infer positions between GPS locations (Bidder et al., 2012; Bidder et al., 2015); combining BLE beacons with other sensors— especially accelerometers—could substantially increase the breadth of questions that could be addressed.

Finally, the current size of BLE beacons (c. 6 g with battery and casing) currently limits deployment to larger species (e.g. over 150 grams in birds). Larger species inherently have larger home ranges (Jenkins, 1981; Lindstedt et al., 1986; Reiss, 1988), making them less prone to positional errors discussed above. However, there is substantial potential for reducing both the footprint and weight of BLE beacons. For example, commercially available beacons are designed to be used with coin batteries (a CR2032 battery), which are both heavy and large. Redesigning beacons to use very small lithium ion batteries could both shed weight and allow a smaller printed circuit board (note that miniaturisation would be limited by the Bluetooth antenna, and small boards may require an external antenna). Commercial BLE beacons are already available that are less than 2 g, and these could be adapted to the Find My network. Doing so would allow beacons to be less invasive on larger animals or to be used with smaller animals, such as smaller bird, mammal, or reptile species, and potentially even on large insects, making miniaturisation therefore a valuable and achievable future direction.

In summary, we have demonstrated that BLE beacons provide an easily accessible tracking solution that can generate data on-par with GPS tracking. We have further provided all necessary information on how to construct, deploy and manage this tracking method, including through custom-designed gateways. Like with all technologies, there are trade-offs, but the light-weight nature of BLE beacons, their low cost, and long battery life should make them particularly useful in studies of animal movements and range use. The potential for high temporal frequency of data collection in areas with high density of Apple devices make them most suited for studies on urban animals, however their use can be extended to natural areas through the deployment of gateways. Further, this simple technology should be readily customisable, allowing the hardware to be miniaturised and combined with other devices and/or on-board sensors. Altogether, BLE beacons should have a wide range of applications, from studying social behaviour, to identifying movement corridors in urban areas through to detecting proximity of animals to roads in reserve areas (Testud et al., 2019). Thus, they will be useful for conservation monitoring and species management as well as ecological studies.

## Author contributions

LMA and DRF conceived the project and the initial design. DRF developed the software and data pipeline. JP, BN and SB undertook field testing, and DRF and SB performed data analysis. All authors contributed to writing the manuscript.

## Acknowledgements

We thank Gerard Borg for helping us with the initial tests of this solution. We also thank Antonio Calatrava for developing and sharing the firmware, Biemster for developing and sharing the interface to the Find My cloud, and to the team who developed the initial openhaystack software that made all of this work possible. The project was conducted with support by the Swiss State Secretariat for Education, Research and Innovation (SERI) under contract number MB22.00056, awarded to LMA. DRF received additional funding from an Eccellenza Professorship Grant of the Swiss National Science Foundation (Grant Number PCEFP3_187058) and grants awarded to DRF from the European Research Council (ERC) under the European Union’s Horizon 2020 research and innovation programme (grant agreement No. 850859). SB and BN were supported by ANU PhD scholarships. All procedures were approved by the ANU Animal Ethics Committee (Projects No. 2022/35, 2023/07), and were conducted under ACT Scientific Licences LT201812 and LT20236.

## Appendices

We provide five appendices with this paper:

*Appendix 1* (generate_keys.py) is the python script that can be used to generate the private and public keys needed for each beacon and to retrieve the data from the cloud.

*Appendix 2* (instructions.txt) is the suggested structure allowing Bluetooth beacon deployments to be managed across multiple projects.

*Appendix 3* (Build_tags_instructions.pdf) contains the instructions for connecting the beacons and flashing the firmware containing the public key.

*Appendix 4* (request_reports_project.py) is a python script used to call down the data from the cloud. This script works with a set folder structure (see Appendix 2) and can be run periodically to append the data into a log file for each project.

*Appendix 5* (BLE_gateway_v4.ino) is the code for using a Freenove ESP32 S3 WROOM development board as a gateway for logging beacon data (also requires a DS3231 real time clock connected to the board). WiFi access (for setting the clock) is controlled via WiFi_name.txt and WiFi_pass.txt files that need to be created on the SD card. Logger ID is provided via a Logger_ID.txt file that needs to be created on the SD card.

