## Supplementary material for "Low-cost animal tracking using Bluetooth low energy beacons on a crowd-sourced network": Data and code: Figure2.R

```
library(lubridate)

load('BLE_test_data.RData')

data$Time <- strptime(data$DateTime,format="%Y-%m-%dT%H:%M:%S")
data$Hour <- hour(data$Time)


get_hist <- function(x) { t <- hour(x)+minute(x)/60; t[t>=23.5] <- t[t>=23.5]-24; a <- hist(t,breaks=seq(-0.5,23.5,1))$counts; return(a) }
a <- lapply(unique(paste(data$Tag.ID,date(data$Time))),function(x) { get_hist(data$Time[which(paste(data$Tag.ID,date(data$Time)) == x)]) })
out <- matrix(unlist(a),ncol=24,byrow=TRUE)
means <- apply(out,2,mean)
maxes <- apply(out,2,max)
mins <- apply(out,2,min)

plot(0:23, means, pch=20, cex=2, xlab="hour (24 hour clock)", ylab="detection rate (detections per hour)", ylim=c(0,35))
arrows(0:23,mins,0:23,maxes,length=0.1,angle=90,code=0)
points(0:23,maxes,pch=20,cex=1)
points(0:23,mins,pch=20,cex=1)
```
