## Supplementary material for "Low-cost animal tracking using Bluetooth low energy beacons on a crowd-sourced network": Data and code: Figure3.R

```
load('BLE_test_data.RData')


library(sp)

latlon.to.UTM <- function(xs, ys) {

	# convert back to lat long for plotting
	data <- data.frame(lon=xs, lat=ys)
	data <- data[!is.na(data$lon),]
	projectionWant <- "+proj=utm +zone=37 +ellps=WGS84"
	coordinates(data) <- c("lon",  "lat")
	proj4string(data) <- CRS("+proj=longlat +datum=WGS84")
	
	data2 <- spTransform(data, CRSobj=projectionWant)

}

data2 <- latlon.to.UTM(data$Lon,data$Lat)
data$Lon.UTM <- data2$lon
data$Lat.UTM <- data2$lat

data2 <- latlon.to.UTM(data$real.Lon,data$real.Lat)
data$real.Lon.UTM <- data2$lon
data$real.Lat.UTM <- data2$lat

data$dist <- sqrt((data$Lon.UTM - data$real.Lon.UTM)^2 + (data$Lat.UTM - data$real.Lat.UTM)^2)
data$log10.dist <- log(data$dist,10)
data$Accuracy2 <- data$Accuracy^2


par(mfrow=c(1,2))


# overview + changepoint FIGURE 3A
library(segmented)
fit_lm = lm(log10.dist ~ 1 + Accuracy, data = data)  # intercept-only model
fit_segmented = segmented(fit_lm, seg.Z = ~Accuracy, npsi = 1)  # Two change points along x
plot(data$Accuracy,data$log10.dist,pch=20,cex=0.5, axes=FALSE, xlab="reported accuracy estimate (m)", ylab="true positional error (log m)",ylim=c(0,4))
axis(1)
axis(2, at=0:4, label=10^(0:4))
box()
plot(fit_segmented,add=T, col="red",lwd=3)
points.segmented(fit_segmented,col="red")


# prob of getting > x m error  FIGURE 3B

data$x.bin <- round(data$Accuracy/10)*10

x <- unique(data$x.bin)
x <- x[order(x)]
plot(NULL, xlim=range(x), ylim=c(0,1), xlab="reported accuracy estimate threshold (m) ", ylab="P(true error > E)")
cols <- colorRampPalette(c("orange","red"))(4)
Es <- c(50,75,100,125)
for (E in 1:length(Es)) {
	y <- sapply(by(data$dist,data$x.bin,function(x) mean(x>Es[E])),identity)
	mod <- glm(y ~ x, family=binomial)
	lines(x,predict(mod, newdata=list(x=x), type="response"), col=cols[E], lwd=2)
}
legend("bottomright", lwd=2, lty=1, legend=Es, col=cols, title="E (m)")

mtext("A", 3, outer=TRUE, at=0.01, line=-3, cex=1.5)
mtext("B", 3, outer=TRUE, at=0.51, line=-3, cex=1.5)
```
