## Supplementary material for "Low-cost animal tracking using Bluetooth low energy beacons on a crowd-sourced network": Data and code: Figure4.r

```
library(lubridate)
library(fields)
library(Rfast)

load('BLE_test_data.RData')

data$Time <- strptime(data$DateTime,format="%Y-%m-%dT%H:%M:%S")
data$time_sec <- difftime(min(data$Time), data$Time, units="secs")

time.thresh <- 300

par(mfrow=c(1,2))

# Pair 1 (road)
Accuracy_cutoffs <- seq(50,120,10)
dists.thresh <- c(50,75,100,125,150,175,200)

props <- matrix(NA, nrow=length(Accuracy_cutoffs), ncol=length(dists.thresh))

for (i in 1:length(Accuracy_cutoffs)) {
	data.1 <- data[which(data$Tag.ID %in% c("A03") & data$Accuracy < Accuracy_cutoffs[i]),]
	data.2 <- data[which(data$Tag.ID %in% c("A04") & data$Accuracy < Accuracy_cutoffs[i]),]


	a <- which(abs(dista(data.1$time_sec,data.2$time_sec)) < time.thresh, arr.ind=TRUE)
	b <- rdist.earth(cbind(data.1$Lon,data.1$Lat),cbind(data.2$Lon,data.2$Lat), miles=FALSE, R= 6371)[a]*1000

	for (j in 1:length(dists.thresh)) {
		together <- sum(!duplicated(a[which(b<dists.thresh[j]),2]) & !duplicated(a[which(b<dists.thresh[j]),1]))
		apart <- sum(!duplicated(a[which(!(b<dists.thresh[j])),2]) & !duplicated(a[which(!(b<dists.thresh[j])),1]))

		props[i,j] <- together/(together+apart)
	}
	
}

cols <- colorRampPalette(c("orange","red"))(length(dists.thresh))
plot(NULL, xlim=range(Accuracy_cutoffs), ylim=c(0,1), xlab="reported accuracy threshold (m)", ylab="P(together within 300 s)")
for (i in 1:length(dists.thresh)) {
	lines(Accuracy_cutoffs,props[,i],col=cols[i], lwd=2)
}
legend("bottomright", lwd=2, col=cols, lty=1, legend=dists.thresh, title="distance threshold (m)", bg="white", cex=0.75)


# Pair 2 (footpath)

Accuracy_cutoffs <- seq(50,120,10)
dists.thresh <- c(50,75,100,125,150,175,200)

props <- matrix(NA, nrow=length(Accuracy_cutoffs), ncol=length(dists.thresh))


data$Time <- strptime(data$DateTime,format="%Y-%m-%dT%H:%M:%S")

for (i in 1:length(Accuracy_cutoffs)) {
	data.1 <- data[which(data$Tag.ID %in% c("A01") & data$Accuracy < Accuracy_cutoffs[i]),]
	data.2 <- data[which(data$Tag.ID %in% c("A02") & data$Accuracy < Accuracy_cutoffs[i]),]

	a <- which(abs(dista(data.1$time_sec,data.2$time_sec)) < time.thresh, arr.ind=TRUE)
	b <- rdist.earth(cbind(data.1$Lon,data.1$Lat),cbind(data.2$Lon,data.2$Lat), miles=FALSE, R= 6371)[a]*1000

	for (j in 1:length(dists.thresh)) {
		together <- sum(!duplicated(a[which(b<dists.thresh[j]),2]) & !duplicated(a[which(b<dists.thresh[j]),1]))
		apart <- sum(!duplicated(a[which(!(b<dists.thresh[j])),2]) & !duplicated(a[which(!(b<dists.thresh[j])),1]))

		props[i,j] <- together/(together+apart)
	}
	
}

cols <- colorRampPalette(c("orange","red"))(length(dists.thresh))
plot(NULL, xlim=range(Accuracy_cutoffs), ylim=c(0,1), xlab="reported accuracy threshold (m)", ylab="P(together within 300 s)")
for (i in 1:length(dists.thresh)) {
	lines(Accuracy_cutoffs,props[,i],col=cols[i], lwd=2)
}
legend("bottomright", lwd=2, col=cols, lty=1, legend=dists.thresh, title="distance threshold (m)", bg="white", cex=0.75)


mtext("A", 3, outer=TRUE, at=0.01, line=-3, cex=1.5)
mtext("B", 3, outer=TRUE, at=0.51, line=-3, cex=1.5)
```
