## Supplementary material for "Low-cost animal tracking using Bluetooth low energy beacons on a crowd-sourced network": Data and code: Figure5_6.R

```
### START FROM HERE
load("Canberra_data_publish.RData")


library(rosm)
library(sp)
library(prettymapr)


utm.to.latlon <- function(lon, lat) {
	require(sp)
	# convert back to lat long for plotting
	data <- data.frame(lon=lon, lat=lat)
	data <- data[!is.na(data$lon),]
	projectionWant <- "+proj=longlat +datum=WGS84"
	coordinates(data) <- c("lon",  "lat")
	proj4string(data) <- CRS("+proj=utm +zone=55 +south=T +ellps=WGS84")
	
	data2 <- spTransform(data, CRSobj=projectionWant)

}

add.alpha <- function(col, alpha=1){
	  if(missing(col))
	    stop("Please provide a vector of colours.")
	  apply(sapply(col, col2rgb)/255, 2, 
                     function(x) 
	                       rgb(x[1], x[2], x[3], alpha=alpha))  
}


## Make 250m squares

tag_data$xs <- round(tag_data$East_UTM/250)*250 - 125
tag_data$ys <- round(tag_data$North_UTM/250)*250 - 125


## Overall profile

intervals <- table(tag_data$timegap[tag_data$same])
intervals <- data.frame(gap=as.numeric(names(intervals)), count=as.numeric(intervals))
intervals$count[seq(2,nrow(intervals),2)] <- intervals$count[seq(2,nrow(intervals),2)] + intervals$count[seq(1,nrow(intervals)-1,2)]
intervals <- intervals[seq(2,nrow(intervals),2),]

days <- length(unique(tag_data$Date[tag_data$same]))
plot(intervals$gap[1:150], intervals$count[1:150]/days, pch=20, xlab="gap between detections (s)", ylab="count", ylim=c(0,50))

# fit a curve
library(aomisc)
intervals$gap.60 <- intervals$gap/60
model <- drm(count/days ~ gap.60, fct = LL.3(),
             data = intervals) ## Log logistic function

#lines(intervals$gap,predict(model), col="red", lwd=2)
abline(v=median(tag_data$timegap[tag_data$same],na.rm=T),lty=2)


week <- TRUE

intervals <- table(tag_data$timegap[tag_data$same & tag_data$week == week])
intervals <- data.frame(gap=as.numeric(names(intervals)), count=as.numeric(intervals))
intervals$count[seq(2,nrow(intervals),2)] <- intervals$count[seq(2,nrow(intervals),2)] + intervals$count[seq(1,nrow(intervals)-1,2)]
intervals <- intervals[seq(2,nrow(intervals),2),]
days <- length(unique(tag_data$Date[tag_data$same& tag_data$week == week]))

# fit a curve
library(aomisc)
intervals$gap.60 <- intervals$gap/60
model <- drm(count/days ~ gap.60, fct = LL.3(),
             data = intervals) ## Log logistic function

lines(intervals$gap,predict(model), col="orange", lwd=2)


week <- FALSE

intervals <- table(tag_data$timegap[tag_data$same& tag_data$week == week])
intervals <- data.frame(gap=as.numeric(names(intervals)), count=as.numeric(intervals))
intervals$count[seq(2,nrow(intervals),2)] <- intervals$count[seq(2,nrow(intervals),2)] + intervals$count[seq(1,nrow(intervals)-1,2)]
intervals <- intervals[seq(2,nrow(intervals),2),]
days <- length(unique(tag_data$Date[tag_data$same& tag_data$week == week]))

# fit a curve
library(aomisc)
intervals$gap.60 <- intervals$gap/60
model <- drm(count/days ~ gap.60, fct = LL.3(),
             data = intervals) ## Log logistic function

lines(intervals$gap,predict(model), col="yellow", lwd=2)

legend("topright", lwd=c(2,2), col=c("orange","yellow"), legend=c("weekdays", "weekends"))


## get spatial profiles


week <- FALSE

redetect <- sapply(by(tag_data$timegap[tag_data$same & tag_data$week == week], paste(tag_data$xs[tag_data$same & tag_data$week == week], tag_data$ys[tag_data$same & tag_data$week == week]), median, na.rm=T),identity)
detect <- sapply(by(tag_data$timegap[tag_data$same & tag_data$week == week], paste(tag_data$xs[tag_data$same & tag_data$week == week], tag_data$ys[tag_data$same & tag_data$week == week]), length),identity)

# remove rarely observed cells (10 observations)
redetect <- redetect[which(detect >= 10)]
detect <- detect[which(detect >= 10)]

# grid
xs <- seq(min(tag_data$xs), max(tag_data$xs), 250)
ys <- seq(min(tag_data$ys), max(tag_data$ys), 250)

grid <- expand.grid(xs=xs,ys=ys)
grid$detect <- detect[match(paste(grid$xs,grid$ys),names(detect))]
grid$redetect <- redetect[match(paste(grid$xs,grid$ys),names(detect))]

grid$xs.lower <- grid$xs-125
grid$xs.upper <- grid$xs+125
grid$ys.lower <- grid$ys-125
grid$ys.upper <- grid$ys+125

d <- utm.to.latlon(grid$xs.lower,grid$ys.lower)
grid$lon.lower <- d$lon
grid$lat.lower <- d$lat
d <- utm.to.latlon(grid$xs.upper,grid$ys.upper)
grid$lon.upper <- d$lon
grid$lat.upper <- d$lat

grid$log.detect <- round(log(grid$detect))
grid$log.redetect <- round(grid$redetect/30)
grid$col <- colorRampPalette(rev(c("#FFFFFF","#FFFF66","#FFFF00","#FF0000")),alpha=FALSE)(20)[round(grid$log.redetect)+1]

### plot above
e <- cbind(c(149.1,149.2),c(-35.35,-35.22))
prettymap(bmaps.plot(bbox(e), project=FALSE, stoponlargerequest=FALSE),scale.style="ticks", scale.linecol="white",scale.label.col="white",drawarrow=TRUE,arrow.scale=0.7,arrow.cols=c("white","white"),arrow.border="white",arrow.text.col="white",arrow.pos="topleft")

for (i in 1:nrow(grid)) {

	if (is.na(grid$col[i])) {
		#rect(grid$lon.lower[i], grid$lat.lower[i], grid$lon.upper[i], grid$lat.upper[i], border=NA,lwd=0.01)
	} else {
		rect(grid$lon.lower[i], grid$lat.lower[i], grid$lon.upper[i], grid$lat.upper[i], col=add.alpha(grid$col[i],0.7),border=NA,lwd=0.01)
	}

}


times1 <- seq(0,max(round(grid$log.redetect),na.rm=T),1)*30
times2 <- seq(0,max(round(grid$log.redetect),na.rm=T),1)*30+30
cols <- colorRampPalette(rev(c("#FFFFFF","#FFFF66","#FFFF00","#FF0000")),alpha=FALSE)(20)[1:(max(round(grid$log.redetect),na.rm=T)+1)]

spaces <- seq(0.001,0.018,0.001)
for (i in 1:length(spaces)) {
	rect(149.2,-35.22-spaces[i]-(i-1)*0.002,149.2-0.0025,-35.22-spaces[i]-0.002-(i-1)*0.002,col=cols[i])
	text(paste(times1[i]," - ",times2[i]," (s)",sep=""),x=149.2-0.00125,y=-35.22-spaces[i]-0.001-(i-1)*0.002, col="white", pos = 4)
}

median(grid$redetect,na.rm=T)
range(grid$redetect,na.rm=T)
```
