## Appendices 1 to 5 for "Low-cost animal tracking using Bluetooth low energy beacons on a crowd-sourced network": generate_keys.py

```
### Code to generate a key for each bluetooth beacon
#
# This code was initially published at https://github.com/biemster/FindMy 
#
# The original code was modified to allow multiple keys to be generated at once
# and for the translated (hex) version of the public keys to be added to a
# text file to ease copying it into the firmware
#
# usage example (in a mac terminal):
# damienfarine$ python generate_keys.py -n 10 -p A -s 0
# 
# this would generate 10 keys, starting with A, and number 0, e.g.:
# A00.keys, A01.keys, ... , A09.keys
#
# it would also produce a file called public_keys.text that contains
# the hexadecimal version of each key (needed for the firmware).
#
###

#!/usr/bin/env python3
import sys,base64,hashlib,random
from cryptography.hazmat.primitives.asymmetric import ec
from cryptography.hazmat.backends import default_backend
import argparse

def int_to_bytes(n, length, endianess='big'):
    return int.to_bytes(n, length, endianess)

def sha256(data):
    digest = hashlib.new("sha256")
    digest.update(data)
    return digest.digest()


parser = argparse.ArgumentParser()
parser.add_argument('-n','--nkeys', help='number of keys to generate', type=int, default=1)
parser.add_argument('-p','--prefix', help='prefix of the keyfiles')
parser.add_argument('-s','--start', help='starting integer for filenames')
args = parser.parse_args()

if args.start:
    s = int(args.start)
else:
    s = 0

for i in range(args.nkeys):

    ok = 0
    while (ok == 0):
        priv = random.getrandbits(224)
        adv = ec.derive_private_key(priv, ec.SECP224R1(), default_backend()).public_key().public_numbers().x
        priv_bytes = int_to_bytes(priv, 28)
        adv_bytes = int_to_bytes(adv, 28)
        priv_b64 = base64.b64encode(priv_bytes).decode("ascii")
        adv_b64 = base64.b64encode(adv_bytes).decode("ascii")
        s256_b64 = base64.b64encode(sha256(adv_bytes)).decode("ascii")

        if '/' in s256_b64[:7]:
            ok = 0
        else:
            ok = 1

    if '/' in s256_b64[:7]:
        print('no key file written, there was a / in the b64 of the hashed pubkey :(')
    else:
        if args.prefix:
            fname = '%s%02d.keys' % (args.prefix, i+s)
        else:
            fname = '%s.keys' % s256_b64[:7]

        with open(fname, 'w') as f:
            f.write('Private key: %s\n' % priv_b64)
            f.write('Advertisement key: %s\n' % adv_b64)
            f.write('Hashed adv key: %s\n' % s256_b64)

        key_hex = base64.b64decode(adv_b64).hex()
        with open('public_keys.txt', 'a') as f:
            f.write(fname + ':\t' + ','.join("0x"+key_hex[i:i + 2] for i in range(0, len(key_hex), 2)) + '\n')
```
