## Appendices 1 to 5 for "Low-cost animal tracking using Bluetooth low energy beacons on a crowd-sourced network": Instructions.txt

Building the folder structure:
Create a base folder (e.g. Bluetooth_deployment) with three subfolders:
./Master_code
./Master_keys
./Projects

Within the ./Projects folder, create a folder for each project, and within this project create two subfolders, e.g.:
./Projects/My_project/Tags
./Projects/My_project/Logs

Within the ./Master_code version, place the 'request_reports_projects.py' file.

Within the ./Master_keys folder, place all of the .keys files generated.

How do the system work?
The request_reports_projects.py script can be called (e.g. from crontab). It is designed to go through each project folder and queries the FindMy database for the data corresponding to the tags who's information is contained in the .keys file in the 'Tags' folder. At present, it is coded to collate data from the past hour (and thus to be run every hour). It then  writes the outputs into the 'Logs' folder. The system will produce one file per day for that project, with the log file containing all of the data from the tags that correspond to the .keys files in the 'Tags' folder.

How do beacons work?
Each bluetooth beacon (tag) is programmed with a unique public key. This key is broadcast by the bluetooth beacon. When this is detected by a phone, the phone logs the key and it's (i.e. the phone's) location from the GPS, and uploads these two bits of information (together with some measure of accuracy) to the FindMy network. When the system queries the FindMy network, it does so by requesting the data using the tag's key.

What are .keys files?
The .keys files contain the relevant keys for each tag to allow the system to download the data from the FindMy network. As well as using the content of these .keys files to query the system, the system uses the filename as the tag name. Thus, if data is coming in for A00.keys, then the log file will contain A00 as the tag name.

Managing deployments
For each project, copy the relevant .keys file into the Tags folder from the Master_keys folder. As the system will download data every hour for the keys with a current file in the Tags folder, this means that undeployed tags should not have a .keys file in the Tags folder until it has been deployed (similarly, the .keys file should be removed if the tag has been removed/fallen off). This will save with data cleaning afterwards (but you should still keep careful records of on/off times).

Naming birds
One nice feature of the system is that tags can be named by simply renaming the relevant .keys file. Say that tag A00 is deployed to 'Alex', one could copy A00.keys to my project's 'Tag' folder, and rename it 'Alex.keys'. Then, all the data would label this tag's data as Alex. This is fine, but requires a little bit of management to avoid losing information about which tag is which. Thus, it is easiest to also keep the A00 in the filename, e.g. 'Alex_A00.keys'. This will return the name Alex_A00, but the _A00 can easily be trimmed out later with a simple R script.

What else?
That's it! Easy.

PLEASE REMEMBER:
(1) Only ever COPY the .keys file from the Master_keys folder, so that if others re-deploy the same tag then they can get a clean copy of this file.
(2) Try to only have the .keys file in the Tag folder when it is actually deployed.
(3) The downloading will only happen if the request_reports_projects.py is run periodically (this can be automated using, for example, the crontab).
