## Appendices 1 to 5 for "Low-cost animal tracking using Bluetooth low energy beacons on a crowd-sourced network": Build_tags_instructions.pdf

SIMPLE INSTRUCTIONS (INSTALL BELOW)

1. **Generate keys using the generate_keys script, e.g.:**

python generate_keys.py -p A -n 100

where A is the starting identifier for the keys files and n is the number of keys to make. This will make files incremental (e.g. A00.keys, A01.keys, etc.). One can also set a starting integer using -s 10 (e.g. to start at A10.keys).

**This produces the file "public_keys.txt" that contains the 28 byte hex key for each .keys file.**

2. **Update the firmware for the current bluetooth beacon:**

Before flashing a bluetooth beacon, we first need to copy the relevant 28 byte hex key corresponding to the the .keys file of interest into the main.c file of openhaystack-alternative (openhaystack-firmware -> apps -> openhaystack-alternative -> main.c), e.g.:

static char public_key[28] = {0x83,0xc3,0x42,0x4d,0xf2,0xef,0x38,0xb4,0x39,0x16,0x10,0x49,0xe5,0x53,0xda,0x3e,0x51,0xd1,0xb7,0x9c,0xc6,0x57,0x63,0x56,0x82,0x42,0x1d,0x5e};

3. **Build the firmware for the tag.**

Open a terminal into the openhaystack-firmware -> apps -> openhaystack-alternative folder, and type:

NRF_MODEL=nrf51 make build

This creates the firmware ready to flash onto the board.

4. **Connect the board to a programmer.**

Next it is required to connect the board to a computer via a programmer. This requires soldering the correct terminals on the programmer to the correct locations on the beacon (see the images below).

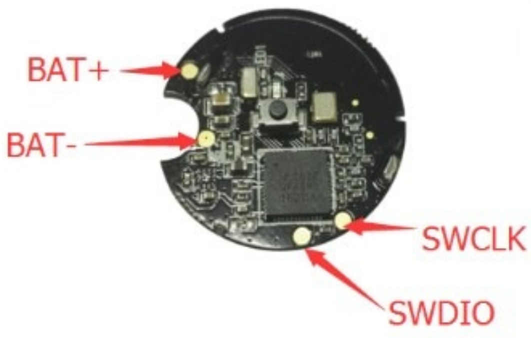


| Pin Number | Pin Name | Pin Function |
|---|---|---|
| 1 | GND | Ground |
| 2 | VDD | 3.3V |
| 3 | SWDIO | Digital pin module data |
| 4 | SWCLK | Digital pin module clock |

5. **Flash the board.**

To flash, in Terminal, run:

openocd -f /usr/local/share/openocd/scripts/interface/stlink.cfg  -f
/usr/local/share/openocd/scripts/target/nrf51.cfg

This will open the connection to the board. Then in a new terminal window, type:

telnet localhost 4444

This will open up a telnet session into the tag. Within this telnet session, type:

```
halt
nrf51 mass_erase

program
/Users/damienfarine/Library/CloudStorage/Dropbox/Bluetooth_beacon/openhaystack-firmware/apps/openhaystack-alternative/compiled/nrf51_firmware.bin verify

program
/Users/damienfarine/Library/CloudStorage/Dropbox/Bluetooth_beacon/openhaystack-firmware/apps/openhaystack-alternative/compiled/nrf51_firmware.bin

resume
```

Then exit from telnet session (type exit) and from the programmer (ctrl+c). Should be done!

**Installation instructions (before the above can be run)**

clone https://github.com/acalatrava/openhaystack-firmware and follow instructions in readme for other installs

```
cd into openhaystack-firmware/apps/openhaystack-alternative
```

Install instructions from the page above:

Get submodules

```
git submodule init
git submodule update
```

Install required dependencies

nRF command line tools

```
brew tap homebrew/cask-drivers
brew install --cask nordic-nrf-command-line-tools
```

binutils

```
brew install binutils
```

gcc-arm-none-eabi

```
brew install --cask gcc-arm-embedded
```
