## Appendices 1 to 5 for "Low-cost animal tracking using Bluetooth low energy beacons on a crowd-sourced network": request_reports_projects.py

```
## 
# Code developed based on https://github.com/biemster/FindMy
# with added capacity to download accuracy data 
# and to work with project-level deployments
# saves data into daily log files within each project
# see Appendix 2 for details on project structure.
#
# Code can be run using:
# python request_reports_projects.py -H 1 -P -B /../Projects
# where 
# -H is the number of hours to go back to look for data
# -P is print to screen (otherwise outputs get appended to a log file)
# -B is the base of the Projects folder
##

#!/usr/bin/env python3
import os,glob
import datetime, time
import argparse,getpass
import base64
import hashlib,hmac
import codecs,struct
import csv
from cryptography.hazmat.primitives import hashes
from cryptography.hazmat.primitives.kdf.pbkdf2 import PBKDF2HMAC
from cryptography.hazmat.primitives.ciphers import Cipher, algorithms, modes
from cryptography.hazmat.primitives.padding import PKCS7
from cryptography.hazmat.primitives.asymmetric import ec
from cryptography.hazmat.backends import default_backend
import objc
from Foundation import NSBundle, NSClassFromString, NSData, NSPropertyListSerialization

def bytes_to_int(b):
    return int(codecs.encode(b, 'hex'), 16)

def sha256(data):
    digest = hashlib.new("sha256")
    digest.update(data)
    return digest.digest()

def decrypt(enc_data, algorithm_dkey, mode):
    decryptor = Cipher(algorithm_dkey, mode, default_backend()).decryptor()
    return decryptor.update(enc_data) + decryptor.finalize()

def unpad(paddedBinary, blocksize):
    unpadder = PKCS7(blocksize).unpadder()
    return unpadder.update(paddedBinary) + unpadder.finalize()

def decode_tag(data):
    latitude = struct.unpack(">i", data[0:4])[0] / 10000000.0
    longitude = struct.unpack(">i", data[4:8])[0] / 10000000.0
    confidence = bytes_to_int(data[8:9])
    status = bytes_to_int(data[9:10])
    return {'lat': latitude, 'lon': longitude, 'conf': confidence, 'status':status}

def readKeychain():
    # https://github.com/libyal/dtformats/blob/main/documentation/MacOS%20keychain%20database%20file%20format.asciidoc
    res = [None] *7
    with open("%s/Library/Keychains/login.keychain-db" % os.path.expanduser("~"),'rb') as db:
        kc = db.read()
        def get_table_offsets(tbl_array_offset):
            ntables = bytes_to_int(kc[tbl_array_offset +4 : tbl_array_offset +8])
            tbl_offsets_b = kc[tbl_array_offset +8 : tbl_array_offset +8 +(ntables *4)]
            return [bytes_to_int(tbl_offsets_b[i:i+4]) +tbl_array_offset for i in range(0, len(tbl_offsets_b), 4)]
        def get_record_offsets(tbl_start):
            nrecords = bytes_to_int(kc[tbl_start +24 : tbl_start +28])
            rec_offsets_b = kc[tbl_start +28 : tbl_start +28 +(nrecords *4)]
            rec_offsets = [bytes_to_int(rec_offsets_b[i:i+4]) +tbl_start for i in range(0, len(rec_offsets_b), 4)]
            return [ro for ro in rec_offsets if ro != tbl_start and bytes_to_int(kc[ro : ro +4])] # remove 0 offset records and empty records
        def match_record_attribute(rec_start, rec_nattr, rec_attr, attr_match):
            attr_offsets_b = kc[rec_start +24 : rec_start +24 +(rec_nattr *4)]
            attr_offsets = [bytes_to_int(attr_offsets_b[i:i+4]) +rec_start -1 for i in range(0, len(attr_offsets_b), 4)]
            if attr_offsets[0] and attr_offsets[0] < rec_start +bytes_to_int(kc[rec_start : rec_start +4]): # non-zero offset, and no weird big values
                if kc[attr_offsets[rec_attr] +4 : attr_offsets[rec_attr] +4 +bytes_to_int(kc[attr_offsets[rec_attr] : attr_offsets[rec_attr] +4])] == attr_match:
                    return kc[rec_start +24 +(rec_nattr *4) : rec_start +24 +(rec_nattr *4) +bytes_to_int(kc[rec_start +16 : rec_start +20])] # return record blob data (NOTE not sure about BLOB size!!!)
            return None
        if kc[:4] == b'kych':
            tbl_offsets = get_table_offsets(bytes_to_int(kc[12:16]))
            symmetric_key_idx = None
            for tbl_start in tbl_offsets[::-1]: # walk backwards so we get the generic password blob before the symmetric key, we need that to select which key to take
                if kc[tbl_start +4 : tbl_start +8] == b'\x00\x00\x00\x11': # Symmetric key
                    rec_offsets = get_record_offsets(tbl_start)
                    for rec_start in rec_offsets:
                        symmetric_key_blob = match_record_attribute(rec_start, 27, 1, symmetric_key_idx) # might be wrong about amount of attributes
                        if symmetric_key_blob:
                            start_crypto_blob = bytes_to_int(symmetric_key_blob[8:12])
                            total_length = bytes_to_int(symmetric_key_blob[12:16])
                            res[2] = symmetric_key_blob[16:24]
                            res[3] = symmetric_key_blob[24 +(start_crypto_blob -0x18) : 24 +(total_length -0x18) ]
                            break
                elif kc[tbl_start +4 : tbl_start +8] == b'\x80\x00\x00\x00': # Generic passwords
                    rec_offsets = get_record_offsets(tbl_start)
                    for rec_start in rec_offsets:
                        icloud_key_blob = match_record_attribute(rec_start, 16, 14, b'iCloud')  # generic password record has 16 attributes
                        if icloud_key_blob:
                            symmetric_key_idx = icloud_key_blob[:20]
                            res[0] = icloud_key_blob[20:28]
                            res[1] = icloud_key_blob[28:]
                            break
                elif kc[tbl_start +4 : tbl_start +8] == b'\x80\x00\x80\x00': # Metadata, containing master key and db key
                    rec_start = get_record_offsets(tbl_start)[0]
                    db_key_blob = kc[rec_start +24 : rec_start +24 +bytes_to_int(kc[rec_start +16 : rec_start +20])] # 2nd record is the one we want
                    res[4] = db_key_blob[44:64]
                    res[5] = db_key_blob[64:72]
                    res[6] = db_key_blob[120:168]
    return res

def retrieveICloudKey():
    icloud_key_IV, icloud_key_enc, symmetric_key_IV, symmetric_key_enc, db_key_salt, db_key_IV, db_key_enc = readKeychain()
    password = getpass.getpass('Keychain password:')
    master_key = PBKDF2HMAC(algorithm=hashes.SHA1(), length=24, salt=db_key_salt, iterations=1000, backend=default_backend()).derive(bytes(password, encoding='ascii'))
    db_key = unpad(decrypt(db_key_enc, algorithms.TripleDES(master_key), modes.CBC(db_key_IV)), algorithms.TripleDES.block_size)[:24]
    p1 = unpad(decrypt(symmetric_key_enc, algorithms.TripleDES(db_key), modes.CBC(b'J\xdd\xa2,y\xe8!\x05')), algorithms.TripleDES.block_size)
    symmetric_key = unpad(decrypt(p1[:32][::-1], algorithms.TripleDES(db_key), modes.CBC(symmetric_key_IV)), algorithms.TripleDES.block_size)[4:]
    icloud_key = unpad(decrypt(icloud_key_enc, algorithms.TripleDES(symmetric_key), modes.CBC(icloud_key_IV)), algorithms.TripleDES.block_size)
    return icloud_key

def getAppleDSIDandSearchPartyToken(iCloudKey):
    # copied from https://github.com/Hsn723/MMeTokenDecrypt
    decryption_key = hmac.new(b't9s\"lx^awe.580Gj%\'ld+0LG<#9xa?>vb)-fkwb92[}', base64.b64decode(iCloudKey), digestmod=hashlib.md5).digest()
    mmeTokenFile = glob.glob("%s/Library/Application Support/iCloud/Accounts/[0-9]*" % os.path.expanduser("~"))[0]
    decryptedBinary = unpad(decrypt(open(mmeTokenFile, 'rb').read(), algorithms.AES(decryption_key), modes.CBC(b'\00' *16)), algorithms.AES.block_size);
    binToPlist = NSData.dataWithBytes_length_(decryptedBinary, len(decryptedBinary))
    tokenPlist = NSPropertyListSerialization.propertyListWithData_options_format_error_(binToPlist, 0, None, None)[0]
    return tokenPlist["appleAccountInfo"]["dsPrsID"], tokenPlist["tokens"]['searchPartyToken']

def getOTPHeaders():
    AOSKitBundle = NSBundle.bundleWithPath_('/System/Library/PrivateFrameworks/AOSKit.framework')
    objc.loadBundleFunctions(AOSKitBundle, globals(), [("retrieveOTPHeadersForDSID", b'')])
    util = NSClassFromString('AOSUtilities')
    anisette = str(util.retrieveOTPHeadersForDSID_("-2")).replace('"', ' ').replace(';', ' ').split()
    return anisette[6], anisette[3]

def getCurrentTimes():
    clientTime = datetime.datetime.utcnow().replace(microsecond=0).isoformat() + 'Z'
    clientTimestamp = int(datetime.datetime.now().strftime('%s'))
    return clientTime, time.tzname[1], clientTimestamp

def writeToFile(ordered, folder):
    date = datetime.datetime.today().strftime("%Y%m%d")
    filename = folder+date+'.csv'
    file_exists = os.path.isfile(filename)
    with open (filename, 'a', encoding='utf8', newline='\n') as csvfile:
        header = ['Tag ID', 'Lat', 'Lon', 'Acc', 'DateTime', 'Link']
        mywriter = csv.writer(csvfile)
        if not file_exists:
            mywriter.writerow(header)
        for rep in ordered: mywriter.writerow([rep['key'],str(rep['lat']),str(rep['lon']),str(rep['conf']),rep['isodatetime'],rep['goog']])


if __name__ == "__main__":
    parser = argparse.ArgumentParser()
    parser.add_argument('-H', '--hours', help='only show reports not older than these hours', type=int, default=24)
    parser.add_argument('-P', '--print', help="option to print direct", action='store_true')
    parser.add_argument('-B', '--base', help="base folder from which to fetch projects")
    args = parser.parse_args()
    iCloud_decryptionkey = retrieveICloudKey()

    AppleDSID,searchPartyToken = getAppleDSIDandSearchPartyToken(iCloud_decryptionkey)
    machineID, oneTimePassword = getOTPHeaders()
    UTCTime, Timezone, unixEpoch = getCurrentTimes()

    request_headers = {
        'Authorization': "Basic %s" % (base64.b64encode((AppleDSID + ':' + searchPartyToken).encode('ascii')).decode('ascii')),
        'X-Apple-I-MD': "%s" % (oneTimePassword),
        'X-Apple-I-MD-RINFO': '17106176',
        'X-Apple-I-MD-M': "%s" % (machineID) ,
        'X-Apple-I-TimeZone': "%s" % (Timezone),
        'X-Apple-I-Client-Time': "%s" % (UTCTime),
        'Content-Type': 'application/json',
        'Accept': 'application/json',
        'X-BA-CLIENT-TIMESTAMP': "%s" % (unixEpoch)
    }

    subfolders = [ f.path for f in os.scandir(args.base) if f.is_dir() ]

    for folder in subfolders:
        if (args.print):
            print('\n\n' + os.path.basename(folder))
        ids = {}
        names = {}
        for keyfile in glob.glob(folder+'/Tags/*.keys'):
            # read key files generated with generate_keys.py
            with open(keyfile) as f:
                hashed_adv = ''
                priv = ''
                name = os.path.basename(keyfile[0:-5])
                for line in f:
                    key = line.rstrip('\n').split(': ')
                    if key[0] == 'Private key':
                        priv = key[1]
                    elif key[0] == 'Hashed adv key':
                        hashed_adv = key[1]
                if priv and hashed_adv:
                    ids[hashed_adv] = priv
                    names[hashed_adv] = name
                else:
                    print("Couldn't find key pair in", keyfile)

        startdate = unixEpoch - 60 * 60 * args.hours
        data = '{"search": [{"endDate": %d, "startDate": %d, "ids": %s}]}' % ((unixEpoch -978307200) *1000000, (startdate -978307200)*1000000, list(ids.keys()))

        # send out the whole thing
        import requests
        data = data.replace("'",'"')
        response = requests.post('https://gateway.icloud.com/acsnservice/fetch', headers=request_headers, data=data)
        #print(response.status_code, response.reason)
        res = response.json()['results']
        #print('%d reports received.' % len(res))

        ordered = []
        found = set()
        for report in res:
            priv = bytes_to_int(base64.b64decode(ids[report['id']]))
            data = base64.b64decode(report['payload'])
            # the following is all copied from https://github.com/hatomist/openhaystack-python, thanks @hatomist!
            timestamp = bytes_to_int(data[0:4])
            if timestamp + 978307200 >= startdate:
                if len(data)>88:
                	data = data[0:4] + data[5:]
                eph_key = ec.EllipticCurvePublicKey.from_encoded_point(ec.SECP224R1(), data[5:62])
                shared_key = ec.derive_private_key(priv, ec.SECP224R1(), default_backend()).exchange(ec.ECDH(), eph_key)
                symmetric_key = sha256(shared_key + b'\x00\x00\x00\x01' + data[5:62])
                decryption_key = symmetric_key[:16]
                iv = symmetric_key[16:]
                enc_data = data[62:72]
                tag = data[72:]
                decrypted = decrypt(enc_data, algorithms.AES(decryption_key), modes.GCM(iv, tag))
                res = decode_tag(decrypted)
                res['timestamp'] = timestamp + 978307200
                res['isodatetime'] = datetime.datetime.fromtimestamp(res['timestamp']).isoformat()
                res['key'] = names[report['id']]
                res['goog'] = 'https://maps.google.com/maps?q=' + str(res['lat']) + ',' + str(res['lon'])
                found.add(res['key'])
                ordered.append(res)

        ordered.sort(key=lambda item: item.get('timestamp'))
        if (args.print):
            print('%d reports used.' % len(ordered))
            for rep in ordered: print(rep['key'], rep['lat'], rep['lon'], rep['conf'], rep['isodatetime'], rep['goog'], sep=',')
            print('found:   ', list(found))
            print('missing: ', [key for key in names.values() if key not in found])
        else:
            writeToFile(ordered,folder+"/Logs/")
```
