## Appendices 1 to 5 for "Low-cost animal tracking using Bluetooth low energy beacons on a crowd-sourced network": BLE_gateway_v4.ino

```
/*
   Based on Neil Kolban example for IDF: https://github.com/nkolban/esp32-snippets/blob/master/cpp_utils/tests/BLE%20Tests/SampleScan.cpp
   Ported to Arduino ESP32 by Evandro Copercini
*/

// sda 8
// scl 9

#include <BLEDevice.h>
#include <BLEUtils.h>
#include <BLEScan.h>
#include <BLEAdvertisedDevice.h>
#include <string.h>
#include "sd_read_write.h"
#include "SD_MMC.h"
#include "WiFi.h"
#include <RTClib.h>
#include "time.h"
#include <math.h>


#define SD_MMC_CMD 38 //Please do not modify it.
#define SD_MMC_CLK 39 //Please do not modify it. 
#define SD_MMC_D0  40 //Please do not modify it.
#define LED_BUILTIN  2
#define uS_TO_S_FACTOR 1000000LL  /* Conversion factor for micro seconds to seconds */


// wifi info for setting the time
char ssid[50];
char password[50];
char logger_id[50];

RTC_DS3231 rtc;

int scanTime = 1; //In seconds
BLEScan* pBLEScan;
int i;
char *knownAddress = "4c00121900"; //Filter string for BLE beacons
char timestamp[20] = "";   //Full date and time stamp
char date[9];           //Short date stamp
int TIME_TO_SLEEP=3;
const int result_length = 10000;
char result[result_length];
char result_tmp[170];
int result_index=0;
char log_result[120];
char filename[55];
size_t cat_buf_len;


// Function that gets time from NTP Server and syncs RTC clock
unsigned long setTime() {

  struct tm timeinfo;
  setenv("TZ", "AEST-10AEDT,M10.1.0,M4.1.0/3",1); // You must include '0' after first designator e.g. GMT0GMT-1, ',1' is true or ON
  getLocalTime(&timeinfo);
  DateTime temp = DateTime(timeinfo.tm_year + 1900, timeinfo.tm_mon + 1, timeinfo.tm_mday, timeinfo.tm_hour, timeinfo.tm_min, timeinfo.tm_sec);
  rtc.adjust(temp);
  return 1;
}

// if it's time to sleep, return 1, else return 0
int update_sleep(int doy, int hour, int minute) {
  
  int dawn = round(328+0.5*152*cos((doy+8-365/2)/58.09));
  int dusk = round(1118-146*cos((doy-365/2)/58.09));
  int current = hour*60 + minute;

  if (current <= dawn) {
    return (dawn - current);
  }
  if (current >= dusk) {
    return(59);
  }
  return 0;
}

int calculateDayOfYear(int day, int month, int year) {
  
  // Given a day, month, and year (4 digit), returns 
  // the day of year. Errors return 999.
  
  int daysInMonth[] = {31,28,31,30,31,30,31,31,30,31,30,31};
  
  // Verify we got a 4-digit year
  if (year < 1000) {
    return 999;
  }
  
  // Check if it is a leap year, this is confusing business
  // See: https://support.microsoft.com/en-us/kb/214019
  if (year%4  == 0) {
    if (year%100 != 0) {
      daysInMonth[1] = 29;
    }
    else {
      if (year%400 == 0) {
        daysInMonth[1] = 29;
      }
    }
   }

  // Make sure we are on a valid day of the month
  if (day < 1) 
  {
    return 999;
  } else if (day > daysInMonth[month-1]) {
    return 999;
  }
  
  int doy = 0;
  for (int i = 0; i < month - 1; i++) {
    doy += daysInMonth[i];
  }
  
  doy += day;
  return doy;
}

int write_to_log(const char *message) {
  DateTime now = rtc.now(); //Read the current date-time from the RTC
  sprintf(timestamp, "%04d-%02d-%02dT%02d:%02d:%02d", now.year(), now.month(), now.day(), now.hour(), now.minute(), now.second());
  sprintf(date, "%04d%02d%02d", now.year(), now.month(), now.day());
  //const size_t cat_buf_len = strlen(timestamp) + 2 + strlen(message) + 4;
  //char* result = (char*) malloc(cat_buf_len);
  //strncpy(result, timestamp, cat_buf_len);
  //snprintf(result, cat_buf_len, "%s%s%s%s", timestamp, " ",message, "\n");
  sprintf(log_result, "%s%s%s%s", timestamp, " ",message, "\n");
  //Serial.printf("writing: %s to log file\n", log_result);
  appendFile(SD_MMC, "/Log.txt", log_result);
  //free(result);

  return 1;
}

class MyAdvertisedDeviceCallbacks: public BLEAdvertisedDeviceCallbacks {
    void onResult(BLEAdvertisedDevice advertisedDevice) {
        //Serial.printf("Advertised Device: %s \n", advertisedDevice.toString().c_str());
        //Serial.printf("Address: %s, Data: %s, rssi: %d\n", advertisedDevice.getAddress().toString().c_str(), manufacturerdata, advertisedDevice.getRSSI());
        if (strstr(advertisedDevice.toString().c_str(), knownAddress) != NULL) {
          char* manufacturerdata = BLEUtils::buildHexData(NULL, (uint8_t*)advertisedDevice.getManufacturerData().data(), advertisedDevice.getManufacturerData().length());
          DateTime now = rtc.now(); //Read the current date-time from the RTC
          sprintf(timestamp, "%04d-%02d-%02dT%02d:%02d:%02d", now.year(), now.month(), now.day(), now.hour(), now.minute(), now.second());
          //sprintf(date, "%04d%02d%02d", now.year(), now.month(), now.day());
          //const size_t cat_buf_len = strlen(timestamp) + 2 + strlen(advertisedDevice.toString().c_str()) + 4;
          //char* result = (char*) malloc(cat_buf_len);
          //strncpy(result, timestamp, cat_buf_len);
          //snprintf(result, cat_buf_len, "%s%s%s%s", timestamp, ",",advertisedDevice.toString().c_str(), "\n");
          //sprintf(result_tmp, "%s%s%s%s", timestamp, ",",advertisedDevice.toString().c_str(), "\n");
          sprintf(result_tmp, "%s,%s,%s,%s,%d\n", timestamp,logger_id,advertisedDevice.getAddress().toString().c_str(), manufacturerdata, advertisedDevice.getRSSI());
          free(manufacturerdata);
          //Serial.printf("%s", result_tmp);
          cat_buf_len = strlen(timestamp) + 2 + strlen(result_tmp) + 4;
          strncat(result, result_tmp, (sizeof(result) - strlen(result)));
          sprintf(result_tmp, "");
          //sprintf(result, "%s%s%s%s", timestamp, ",",advertisedDevice.toString().c_str(), "\n");
          result_index = strlen(result);
          //const size_t cat_buf_len2 = strlen(date) + 6;
          //char* filename = (char*) malloc(cat_buf_len2);
          //strncpy(filename, date, cat_buf_len2);
          //snprintf(filename, cat_buf_len2, "/%s%s", date, ".txt");
          if (result_index > (result_length-120)) {
            sprintf(date, "%04d%02d%02d", now.year(), now.month(), now.day());
            sprintf(filename, "/%s_%s%s", logger_id, date, ".txt");
            appendFile(SD_MMC, filename, result);
            //Serial.printf("%s", result);
            sprintf(result, "");
          }
          
          digitalWrite(LED_BUILTIN, HIGH);   // turn the LED on (HIGH is the voltage level)
          delay(10);                       // wait for a second
          digitalWrite(LED_BUILTIN, LOW);    // turn the LED off by making the voltage LOW
          //free(result); 
          //free(filename); 
          //free(cat_buf_len);
          //free(cat_buf_len2);
        }

    }
};

void print_wakeup_reason(){
  esp_sleep_wakeup_cause_t wakeup_reason;

  wakeup_reason = esp_sleep_get_wakeup_cause();

  switch(wakeup_reason){
    case ESP_SLEEP_WAKEUP_EXT0 : write_to_log("Wakeup caused by external signal using RTC_IO"); break;
    case ESP_SLEEP_WAKEUP_EXT1 : write_to_log("Wakeup caused by external signal using RTC_CNTL"); break;
    case ESP_SLEEP_WAKEUP_TIMER : write_to_log("Wakeup caused by timer"); break;
    case ESP_SLEEP_WAKEUP_TOUCHPAD : write_to_log("Wakeup caused by touchpad"); break;
    case ESP_SLEEP_WAKEUP_ULP : write_to_log("Wakeup caused by ULP program"); break;
    default : write_to_log("Wakeup was not caused by deep sleep"); break;
  }
}


void setup() {
  

  Serial.begin(115200);
  delay(1000);

  Serial.println("\nBooting up");

  // initialize digital pin LED_BUILTIN as an output.
  pinMode(LED_BUILTIN, OUTPUT);

  // set up memory card
  SD_MMC.setPins(SD_MMC_CLK, SD_MMC_CMD, SD_MMC_D0);
  if (!SD_MMC.begin("/sdcard", true, true, SDMMC_FREQ_DEFAULT, 5)) {
    Serial.println("Card Mount Failed");
      while (1) { // flash leds continuously
        digitalWrite(LED_BUILTIN, HIGH);   // turn the LED on (HIGH is the voltage level)
        delay(500);                       // wait for a second
        digitalWrite(LED_BUILTIN, LOW);    // turn the LED off by making the voltage LOW
        delay(200);                       // wait for a second
        digitalWrite(LED_BUILTIN, HIGH);   // turn the LED on (HIGH is the voltage level)
        delay(200);                       // wait for a second
        digitalWrite(LED_BUILTIN, LOW);   // turn the LED on (HIGH is the voltage level)
        delay(200);                       // wait for a second
      }
  }
  uint8_t cardType = SD_MMC.cardType();
  if(cardType == CARD_NONE){
      Serial.println("No SD_MMC card attached");
      digitalWrite(LED_BUILTIN, HIGH);   // turn the LED on (HIGH is the voltage level)
      while (1) { // flash leds continuously
        digitalWrite(LED_BUILTIN, HIGH);   // turn the LED on (HIGH is the voltage level)
        delay(500);                       // wait for a second
        digitalWrite(LED_BUILTIN, LOW);    // turn the LED off by making the voltage LOW
        delay(200);                       // wait for a second
        digitalWrite(LED_BUILTIN, HIGH);   // turn the LED on (HIGH is the voltage level)
        delay(200);                       // wait for a second
        digitalWrite(LED_BUILTIN, LOW);   // turn the LED on (HIGH is the voltage level)
        delay(200);                       // wait for a second
      }
  }

  esp_sleep_enable_timer_wakeup(TIME_TO_SLEEP * uS_TO_S_FACTOR);

  Serial.println("starting RTC module ");
  if (! rtc.begin()) {
    //Serial.println("RTC module is NOT found");
    write_to_log("RTC module is NOT found");
    while (1) { // keep led on continuously
        digitalWrite(LED_BUILTIN, HIGH);   // turn the LED on (HIGH is the voltage level)
    }
  }
  Serial.println("RTC module is found");
  Serial.println("Checking clock time");
  DateTime now = rtc.now(); //Read the current date-time from the RTC

  File file;
  unsigned int index;
  char next_char;
  if (rtc.lostPower() || int(now.year()) < 2024) {
    Serial.println("RTC lost power or incorrect, setting time");
    write_to_log("RTC lost power or incorrect, setting time");


    Serial.println("Reading SSID from file:");
    // read in details from SD
    file = SD_MMC.open("/WiFi_name.txt");
    if (!file) {
      write_to_log("Opening WiFi name file failed");
      return;
    }
    index = 0;
    next_char;
    while (file.available()) {
      next_char = file.read();
      if (next_char != 13 & next_char != '\n' & next_char != 10) ssid[index++] = next_char;
      //ssid[index++] = next_char;
      //String ssid = file.readStringUntil('\r');
      //Serial.println(ssid);
    }
    //ssid[index] = '/0';
    file.close();
    Serial.println(ssid);

    Serial.println("Reading WiFi password from file:");
    // read in details from SD
    file = SD_MMC.open("/WiFi_pass.txt");
    if (!file) {
      write_to_log("Opening WiFi pass file failed");
      return;
    }
    index = 0;
    while (file.available()) {
      next_char = file.read();
      if (next_char != 13 & next_char != '\n' & next_char != 10) password[index++] = next_char;
      //password[index++] = next_char;
      //String password = file.readStringUntil('\r');
    }
    //password[index] = '/0';
    file.close();
    Serial.println(password);

    write_to_log("starting WIFI ");
    Start_WiFi(ssid,password);
    configTime(0, 0, "pool.ntp.org");
    delay(1000);
    setTime();
    delay(500);
  }
  
  // update in case time was reset
  now = rtc.now(); //Read the current date-time from the RTC
  //sprintf(timestamp, "%04d-%02d-%02dT%02d:%02d:%02d", now.year(), now.month(), now.day(), now.hour(), now.minute(), now.second());
  //Serial.println(timestamp);

  if (int(now.year()) < 2024) {
    Serial.println("RTC appears incorrect");
    while (1) { // flash leds continuously
      digitalWrite(LED_BUILTIN, HIGH);   // turn the LED on (HIGH is the voltage level)
      delay(200);                       // wait for a second
      digitalWrite(LED_BUILTIN, LOW);    // turn the LED off by making the voltage LOW
      delay(200);                       // wait for a second
    }
  }

  //setenv("TZ", "AEST-10AEDT,M10.1.0,M4.1.0/3",1);
  setenv("TZ", "AEST-10", 1);
  tzset();

  write_to_log("RTC & SD card OK");

  write_to_log("Reading Logger ID");
  Serial.println("Reading Logger:");

  file = SD_MMC.open("/Logger_ID.txt");
  if (!file) {
    write_to_log("Opening Logger_ID file failed");
    return;
  }
  index = 0;
  while (file.available()) {
    next_char = file.read();
    if (next_char != 13 & next_char != '\n' & next_char != 10) logger_id[index++] = next_char;
  }
  //logger_id[index] = '/0';
  file.close();
  Serial.println(logger_id);

  now = rtc.now(); //Read the current date-time from the RTC
  sprintf(timestamp, "%04d-%02d-%02dT%02d:%02d:%02d", now.year(), now.month(), now.day(), now.hour(), now.minute(), now.second());
  Serial.print("Time is: ");
  Serial.println(timestamp);

  //Print the wakeup reason for ESP32
  print_wakeup_reason();
  
  // switch of WIFI for goo
  WiFi.disconnect();
  WiFi.mode(WIFI_OFF);
  write_to_log("WIFI stopped");


  BLEDevice::init("");
  pBLEScan = BLEDevice::getScan(); //create new scan
  pBLEScan->setAdvertisedDeviceCallbacks(new MyAdvertisedDeviceCallbacks());
  pBLEScan->setActiveScan(true); //active scan uses more power, but get results faster
  pBLEScan->setInterval(100);
  pBLEScan->setWindow(99);  // less or equal setInterval value


  // if all OK, blink 5 times
  for (i = 1; i < 6; ++i)
  {
    digitalWrite(LED_BUILTIN, HIGH);   // turn the LED on (HIGH is the voltage level)
    delay(500);                       // wait for a second
    digitalWrite(LED_BUILTIN, LOW);    // turn the LED off by making the voltage LOW
    delay(500);                       // wait for a second
  }

  write_to_log("Starting logging");
  Serial.println("Starting logging");

}


int Start_WiFi(const char* ssid, const char* password){
 int connAttempts = 0;
 Serial.println("\r\nConnecting to: "+String(ssid));
 WiFi.begin(ssid, password);
 while (WiFi.status() != WL_CONNECTED ) {
   delay(500);
   Serial.print(".");
   if(connAttempts > 20) {
    write_to_log("Failed to connect to WiFi!");
    return -5;
   }
   connAttempts++;
 }
 write_to_log("setting the time!");
 return 1;
}


void loop() {

  for (i = 0; i < 60; ++i)
  {
    BLEScanResults foundDevices = pBLEScan->start(scanTime, false);
    //Serial.print("Devices found: ");
    //Serial.println(foundDevices.getCount());
    //Serial.println("Scan done!");
    pBLEScan->clearResults();   // delete results fromBLEScan buffer to release memory
    delay(100);
  }

  DateTime now = rtc.now(); //Read the current date-time from the RTC

  // write any unsaved results
  if (strlen(result) > 20) {
    sprintf(date, "%04d%02d%02d", now.year(), now.month(), now.day());
    sprintf(filename, "/%s_%s%s", logger_id, date, ".txt");
    appendFile(SD_MMC, filename, result);
    //Serial.printf("Output2\n");
    //Serial.printf("%s\n", result);
    sprintf(result, "");
  }
 
  
  //char buffer1[30];
  //sprintf(buffer1, "Heap size: %d\n", ESP.getFreeHeap());
  //Serial.printf("%s", buffer1);
  //write_to_log(buffer1);
  
  Serial.println("Checking sleep time");
  // if time to sleep, then sleep for 59 minutes
  int doy = calculateDayOfYear(int(now.day()),int(now.month()),int(now.year()));
  int sleep_time = update_sleep(doy,int(now.hour()),int(now.minute()));

  if (sleep_time>0)  {
    if (sleep_time >= 59) {
      TIME_TO_SLEEP = 59*60;
    } else {
      TIME_TO_SLEEP = sleep_time*60;
    }
    char buffer[30];
    sprintf(buffer, "Going to sleep for %d minutes", TIME_TO_SLEEP/60);
    write_to_log(buffer);

    esp_sleep_enable_timer_wakeup(TIME_TO_SLEEP * uS_TO_S_FACTOR);

    delay(1000);
    Serial.flush(); 
    esp_deep_sleep_start();
  }
  
}
```
